# Cosine Similarity Conflates Clinically Distinct Cancer Variants: A Case for Typed-Graph Retrieval in Precision Oncology Decision Support

**DOI:** 10.64898/2026.05.05.723102

**Authors:** Umair Khan

## Abstract

Retrieval-augmented generation (RAG) is increasingly applied to clinical decision support in oncology, where treatment selection depends on identifying a patient’s specific somatic variant from an NGS report and matching it to evidence-graded therapy options. The vector retrieval that underlies most RAG systems uses cosine similarity over text embeddings, an architecture optimized for linguistic proximity rather than entity-level identity. We hypothesize that cosine-similarity-based retrieval conflates clinically distinct cancer variants at clinically relevant rates, while a typed-graph approach in which each variant is a discrete node preserves variant-level identity by construction.

We evaluated 9 cancer variant pairs with differential FDA-approved therapy indications, variant identity informed by the CIViC clinical variant evidence database and primary clinical literature. The pairs span BRAF, EGFR, KRAS, ERBB2, PIK3CA, and NTRK1, including the canonical EGFR L858R vs T790M sensitivity-versus-resistance pair and KRAS G12C vs G12D (only G12C has an FDA-approved targeted therapy). We computed pairwise cosine similarity across three open-source embedding models (PubMedBERT, MedCPT, BGE-large-en-v1.5) and three text formats, and in this revision added template-matched negative controls, formal separation metrics, variant-specific long-format text, corpus-level ranked retrieval, an exact-ID guardrail baseline, and an end-to-end typed-graph pipeline.

Across the medium format (gene + variant + tumor type), **100% of clinically distinct variant pairs (9/9) had cosine similarity ≥ 0.95 under both biomedical encoders** (PubMedBERT, MedCPT; exact binomial 95% CI [66.4%, 100%]); the general-purpose encoder (BGE-large-en-v1.5) conflated 11%. The biomedically pre-trained encoders performed *worse*, not better, than the general-purpose encoder. Template-matched but biologically unrelated negative controls scored lower than the clinically distinct pairs, confirming the high similarities are genuine conflation and not a template artifact, and equivalent-notation positives were not separable from clinically distinct pairs by a cosine threshold (medium-format ROC-AUC ≤ 0.54, below 0.5 for the biomedical encoders). In corpus-level ranked retrieval over a 52-document corpus, the wrong paired variant appeared in the top 5 for 75% to 100% of queries under the biomedical encoders. Adding an exact variant-ID guardrail to cosine retrieval, and routing retrieval through an end-to-end typed-graph pipeline, both reduced wrong-variant retrieval to zero. We argue that, conditional on a normalization layer that resolves variant strings to canonical nodes, typed-graph retrieval, or vector retrieval coupled with strict variant-ID guardrails, should be the default substrate for variant-level clinical decision support. We empirically validate the typed-graph baseline end-to-end on the same nine-pair benchmark, demonstrating 95.8% normalization accuracy and a 0% wrong-variant retrieval rate.

## 1. Introduction

Targeted therapy in oncology depends on identifying the patient’s specific somatic variant. The decision to prescribe osimertinib for non-small-cell lung cancer hinges on which EGFR variant is present and in what clinical context: an L858R substitution at first diagnosis is a sensitizing mutation that makes osimertinib first-line standard of care [1], while T790M emerging at progression on a first-generation EGFR tyrosine kinase inhibitor is a resistance mutation that makes osimertinib the second-line salvage [2]. Two variants in the same gene; opposite clinical implications; one drug indicated in different settings for different reasons. Confusing them is not a small error.

The same pattern recurs across precision oncology. KRAS G12C in NSCLC has two FDA-approved covalent inhibitors (sotorasib [3] and adagrasib [4]) whose mechanism depends on the cysteine residue at position 12. KRAS G12D, the same protein, the same residue, a different amino acid substitution, has no FDA-approved targeted therapy as of this writing — the noncovalent inhibitors that target G12D are entirely different chemotypes [5]. BRAF V600E and V600K are both melanoma drivers that respond to combined BRAF and MEK inhibition, but the V600E patients in the COMBI-d trial had numerically better progression-free survival than V600K patients [6], and the demographic profile of V600K-mutant melanoma differs from V600E [7]. NTRK1 gene fusions are responsive to larotrectinib and entrectinib (tumor-agnostic FDA approval); NTRK1 point mutations are not.

These distinctions are clinically actionable. They drive drug selection. They are also linguistically subtle. The strings “BRAF V600E” and “BRAF V600K” differ by one character. The strings “EGFR L858R” and “EGFR T790M” share three of five tokens. Any retrieval system that operates on text similarity over cancer variant strings is at risk of treating these as equivalent.

This matters because retrieval-augmented generation (RAG) [8,9] has become the default architecture for grounding large language model output in domain-specific knowledge bases, including in clinical applications [10–13]. The standard RAG pipeline embeds both the query and the candidate evidence into a vector space, scores them by cosine similarity, returns the top-k nearest matches, and conditions the LLM’s generation on those matches. The architecture is engineered for linguistic proximity.

For variant-level clinical reasoning, linguistic proximity is the wrong primitive. Two variants either refer to the same molecular event or they don’t. The relationship is binary, identity-based, and discrete. Cosine similarity is continuous and similarity-based. The mismatch between the operation and the data is structural.

A typed knowledge graph, by contrast, represents each variant as a discrete node connected to drugs, trials, evidence statements, and contraindications by typed edges [14,15]. Identity is preserved by construction: there is exactly one node for BRAF V600E and exactly one node for BRAF V600K, with separate outgoing edges to therapy options and evidence references. Retrieval is graph traversal, not similarity scoring. Two variants that differ by a single amino acid are two distinct nodes regardless of how textually close their string representations are.

This paper quantifies the gap. We evaluate the rate at which standard vector retrieval, using widely-deployed biomedical and general-purpose embedding models, conflates clinically distinct cancer variant pairs. We compare to the typed-graph baseline, where conflation is by construction zero. We then argue that the architectural implication for clinical decision support is unambiguous: variant-level retrieval should not run on cosine similarity, even when the embedding model is biomedically pre-trained.

Our contributions are:

1. **A benchmark of variant-pair conflation rates** with exact binomial 95% confidence intervals across three commonly deployed open-source embedding models (PubMedBERT, MedCPT, BGE-large-en-v1.5).
2. **Template-matched unrelated negative controls** quantifying the baseline similarity floor and confirming that the observed high similarities reflect true conflation rather than a template artifact.
3. **Formal separation metrics** (ROC-AUC, Kolmogorov-Smirnov statistic, Cliff’s delta) between equivalent-notation positives and clinically distinct pairs.
4. **A failure-mode characterization** showing that conflation rates are not improved by biomedical pre-training, by larger embedding models, or by additional clinical context, using variant-specific asymmetric long-format text.
5. **Corpus-level ranked retrieval results** demonstrating that pairwise conflation translates to wrong-variant@k errors in production-like ranked retrieval.
6. **An empirical comparison of three architectures**: cosine-only retrieval, cosine retrieval with an exact variant-ID guardrail, and an end-to-end typed-graph pipeline with ingestion-time normalization.
7. **An honest characterization** of where the typed-graph approach’s normalization layer succeeds (point mutations) and where it requires additional engineering (structural variants).
8. **An open benchmark** of the variant-pair set, embedding scripts, retrieval corpus, and result data for replication and extension at *github*.*com/unmirihealth/unmiri-ngs-fhir-schema* (manuscript supplement) and a Zenodo deposit with DOI [16].

We do not claim that vector retrieval has no place in clinical NLP. Literature search across PubMed abstracts, retrieval over unstructured clinical notes, and de-duplication across LIMS imports are all well-suited to dense retrieval. Our argument is narrower: when retrieval is the substrate for selecting a targeted therapy based on a patient’s specific variant, the architecture must preserve variant-level identity. Cosine similarity does not.

## 2. Related Work

### 2.1 Vector retrieval-augmented generation

Retrieval-augmented generation in its modern form was formalized by Lewis et al. [8], with the RAG architecture combining a parametric language model with non-parametric retrieval over a dense passage index. Subsequent work refined the dense-retriever side of the pipeline [9,17] and scaled the retrieval corpus to trillions of tokens [18]. A recent survey by Gao et al. [19] organizes the field into Naive, Advanced, and Modular RAG taxonomies and catalogs documented limitations including retrieval noise, position-in-context effects [20], and grounding failures [21].

Within biomedicine, dense retrievers built on PubMedBERT [22], BioGPT [23], and MedCPT [24] have become the default substrate for biomedical retrieval. Recent benchmarks of medical RAG systems demonstrate that retrieval-side failures persist even with biomedically pre-trained encoders [25]. Almanac [26] is a clinical-RAG system evaluated by board-certified physicians and represents the current state of clinical RAG deployment.

The architectural assumption shared across this literature is that linguistic proximity is a reasonable proxy for relevance. For most retrieval tasks — finding documents that discuss a topic, finding answer-bearing passages for a question, finding analogous clinical cases — this assumption is sound. For variant-level clinical reasoning, it is the assumption we challenge.

### 2.2 Knowledge-graph-augmented retrieval and reasoning

A parallel line of work integrates large language models with structured knowledge graphs. Microsoft’s GraphRAG [14] demonstrates that graph-indexed retrieval outperforms vector RAG on global, structural questions where entity relationships matter more than text similarity. Think-on-Graph [27] shows that beam-search reasoning over typed knowledge-graph edges yields traceable, correctable inference. G-Retriever [15] explicitly addresses hallucination by routing retrieval through graph structure rather than text similarity. Reasoning on Graphs [28] develops the planning-retrieval-reasoning framework for faithful and interpretable LLM reasoning. The recent IEEE TKDE survey by Pan et al. [29] organizes the LLM+KG research program.

In biomedicine specifically, KG-RAG over the SPOKE biomedical knowledge graph [30] is the closest published analogue to the architecture we advocate. PrimeKG [31] is a worked example of a typed precision-medicine KG with genes, variants, drugs, and diseases as distinct typed nodes, preserving entity identity by construction. Large language models with retrieval have also been applied directly to precision-oncology decision support, including molecular-tumor-board recommendation [45].

While cosine-similarity limitations for exact entity matching are recognized in general information retrieval, and graph-RAG architectures for medical safety have been proposed (SPOKE [30], PrimeKG [31], Microsoft GraphRAG [14]), the specific empirical quantification of variant-level conflation in precision oncology, and the counterintuitive finding that biomedical pre-training exacerbates rather than mitigates it, has not previously been systematically benchmarked. This paper makes that targeted empirical contribution; the v2 revision adds template-matched negative controls, separation metrics, corpus-level retrieval, and an end-to-end typed-graph baseline so the architectural recommendation rests on measurement rather than assertion.

Watson for Oncology [32,33] is the most prominent cautionary precedent. Internal documents and post-hoc analyses describe a system that recommended unsafe therapies grounded in retrieval over clinical literature without typed entity-level identity in the substrate. Cross-vendor commercial NGS interpretation tools have similarly struggled with concordance across labs [34], reinforcing that the variant-identity problem is not specific to any one architecture.

### 2.3 Clinical variant interpretation and the AMP/ASCO/CAP framework

Clinical interpretation of cancer variants is governed by professional society guidelines. The 2017 AMP/ASCO/CAP framework [35] is the current standard for somatic variant interpretation in cancer, with a four-tier taxonomy: Tier I (variants of strong clinical significance, with sub-Levels A and B), Tier II (variants of potential clinical significance, with sub-Levels C and D), Tier III (variants of unknown clinical significance), and Tier IV (variants deemed benign or likely benign). Throughout this paper we use the compact form “Tier I-A” as shorthand for “Tier I, Level A” (and similarly for I-B, II-C, II-D); this shorthand is widely used in secondary literature and matches the schema enum values in [16]. The 2015 ACMG/AMP guidelines [36] govern germline variant interpretation. The 2022 ClinGen/CGC/VICC oncogenicity classification [37] separates oncogenicity from clinical actionability. ESMO’s ESCAT framework [38] provides an alternative actionability scale.

Knowledge bases that operationalize these taxonomies include CIViC (Clinical Interpretation of Variants in Cancer) [39], a CC0 public-domain knowledge base built on community curation; ClinVar [40], the NIH-maintained germline variant evidence repository; and COSMIC [41], the Catalogue of Somatic Mutations in Cancer (free for academic use, paid commercial license required for curated content). OncoKB [42] is a precision oncology knowledge base whose therapeutic actionability levels are widely cited (commercial license required for curated content; not used as a data source in this study).

### 2.4 Clinical decision support architecture and the variant-conflation failure mode

Clinical decision support (CDS) systems in oncology face documented failure modes including alert malfunctions [43] and inappropriate recommendations driven by retrieval-side errors. The FDA’s 2022 CDS guidance [44] establishes a regulatory framework that distinguishes Device CDS from Non-Device CDS based on (among other criteria) whether the system surfaces sources transparently and whether the clinician can independently review the basis of the recommendation. A retrieval architecture that conflates variants forecloses transparent source review by design: if the system cannot distinguish V600E from V600K in its substrate, it cannot tell the clinician which variant the recommendation is based on.

Cross-vendor heterogeneity in commercial NGS reports compounds the problem [34]. Foundation Medicine, Tempus, Caris Life Sciences, Guardant Health, and Natera each ship reports with vendor-specific variant nomenclature, biomarker conventions, and therapy-tier vocabularies. Any retrieval system that ingests reports from more than one lab must distinguish variants at the per-vendor level even after normalization, which makes the variant-identity property of the retrieval substrate a load-bearing engineering decision, not a mere implementation detail.

## 3. Methods

### 3.1 Variant pair selection

We curated a set of cancer variant pairs informed by the CIViC clinical variant evidence database [39] and primary clinical literature, where each pair (A, B) satisfies three criteria:

1. Both variants are documented in CIViC (Variant IDs in `variant-pairs.csv`) and supported by primary clinical literature with strong clinical or trial-level evidence.
2. The clinical evidence describes materially different therapeutic indications, including: different FDA-approved drugs, different lines of therapy, sensitivity vs resistance distinctions, or eligibility differences in active clinical trials.
3. The variants are within the same gene to ensure the textual representations differ only at the variant-specifying tokens, isolating the conflation question from broader gene-level disambiguation.

The variant-pair set used in the primary analysis comprises nine pairs covering the most commonly encountered clinically distinct variant relationships in current FDA-approved targeted therapy. CIViC Variant IDs (VIDs) for each variant, where indexed, are provided in `variant-pairs.csv`:

- BRAF V600E (CIViC VID 12) vs BRAF V600K (VID 563), in melanoma
- EGFR L858R (VID 33) vs EGFR T790M (VID 34), in NSCLC; sensitizing vs resistance
- EGFR exon 19 deletion / E746_A750del (VID 1002) vs EGFR L858R (VID 33), in NSCLC; both sensitizing, drug-specific differences
- KRAS G12C (VID 78) vs KRAS G12D (VID 79), in NSCLC; covalent inhibitor target vs no FDA-approved targeted therapy
- KRAS G12C (VID 78) vs KRAS G12V (VID 425), in NSCLC; covalent inhibitor target vs no FDA-approved targeted therapy as of 2025
- ERBB2 amplification (VID 306) vs ERBB2 G776delinsVC (not discretely indexed in CIViC; canonical entry is the categorical “ERBB2 exon 20 insertion” variant), in NSCLC; different therapeutic class
- PIK3CA E545K (VID 104) vs PIK3CA M1043I (VID 937), in breast cancer; different SOLAR-1 / INAVO120 eligibility
- PIK3CA H1047R (VID 107) vs PIK3CA E542K (VID 103), in breast cancer; allele-specific SOLAR-1 hazard ratio differences
- NTRK1 fusion (VID 1278; LMNA::NTRK1 — the most prominently indexed NTRK1-fusion entry) vs NTRK1 point mutation (VID 2690; G595R solvent-front kinase-domain variant), in NSCLC and tumor-agnostic; FDA-targetable vs not currently FDA-targetable in NSCLC

CIViC VIDs were verified against civicdb.org/variants/<VID> URLs. Per-variant Evidence Items (EIDs) are not enumerated in the manuscript because the CIViC v2 data model attaches Evidence Items to Molecular Profiles rather than directly to Variants, and individual EID stability is dependent on ongoing curator activity; readers can retrieve current evidence for any listed variant by querying the CIViC GraphQL API or browsing the variant page.

The full variant-pair set with the metadata above is provided as supplementary material in `variant-pairs.csv`. For each variant we additionally included a redundant-form pair (e.g., “EGFR L858R” vs “EGFR p.L858R”) as a positive control, expected to score ≈ 1.0 cosine similarity, demonstrating that the embedding models do recognize when variants are textually equivalent.

### 3.2 Embedding models

We tested three open-source biomedical and general-purpose text embedding models, all available without API authentication:

1. **PubMedBERT** [22] (`microsoft/BiomedNLP-BiomedBERT-base-uncased-abstract-fulltext`). Encoder pre-trained on PubMed abstracts and PubMedCentral full text.
2. **MedCPT** [24] (`ncbi/MedCPT-Article-Encoder`, `ncbi/MedCPT-Query-Encoder`). Contrastive-pretrained biomedical retriever, trained on PubMed search-log data, designed specifically for biomedical retrieval rather than general embedding.
3. **BGE-large-en-v1.5** (`BAAI/bge-large-en-v1.5`). General-purpose state-of-the-art text embedding, included as a non-biomedical baseline.

We deliberately did not test commercial proprietary embedding APIs (OpenAI text-embedding-3-large, Voyage AI, Cohere) for two reasons. First, replication: closed-API embeddings are rate-limited, costed, and may change without notice. Second, fairness: the strongest argument against vector retrieval is that even biomedically-tuned, open-source state-of-the-art embeddings exhibit conflation. If the open models conflate, the proprietary models are unlikely to materially change the architectural conclusion.

### 3.3 Text formats

For each variant we generated three text representations to test the hypothesis that adding clinical context changes retrieval behavior:

- **Short**: `<gene-symbol> <variant>` (e.g., “BRAF V600E”).
- **Medium**: `<gene-symbol> <variant> in <tumor-type>` (e.g., “BRAF V600E in melanoma”).
- **Long**: `<gene-symbol> <variant> in <tumor-type>`; `<one-sentence-clinical-context>` (e.g., “BRAF V600E in melanoma; targetable with combined BRAF and MEK inhibition (dabrafenib + trametinib).”).

The clinical-context sentence for the long form is drawn from CIViC evidence statements only, with no proprietary content.

### 3.4 Similarity computation

For each variant pair (A, B), each embedding model M, and each text format F, we computed the cosine similarity between the L2-normalized embedding vectors of A and B:

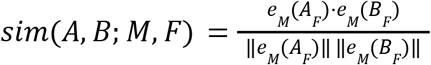

This yields 9 pairs × 3 models × 3 formats = 81 measurements for the clinically distinct pairs, plus an equivalent-form positive control set.

### 3.5 Conflation thresholds

We define **conflation rate** at threshold τ as the proportion of clinically distinct variant pairs whose cosine similarity meets or exceeds τ:

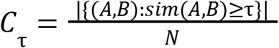

We report rates at τ = 0.85, 0.90, 0.95, and 0.99. The choice of τ is informed by typical top-k retrieval thresholds in deployed RAG systems: similarity ≥ 0.95 is commonly within the top-3 retrieval window, and ≥ 0.99 is commonly the criterion for “near-duplicate” detection.

### 3.6 Typed-graph baseline

The typed-graph baseline represents each variant as a discrete node in a property graph (implemented for this study as an in-memory networkx graph, so that the experiment carries no database dependency; the same schema maps directly onto a Neo4j property graph) with the following node and edge schema:

- Nodes: `Gene, Variant, Drug, Trial, EvidenceStatement, TumorType`.
- Variant nodes carry properties: gene (HGNC), `hgvs_coding, hgvs_protein, variant_type, assembly, genomic_coordinate`.
- Edges: (Variant)-[:INDICATES_RESPONSE_TO {evidence_level, source}]→(Drug), (Variant)-[:CONTRAINDICATES {reason, evidence_level}]→(Drug), (Variant)-[:ENROLLED_IN]→(Trial), (Variant)-[:DESCRIBED_BY]→(EvidenceStatement).

Variant retrieval in the typed-graph baseline is exact match on the (gene, hgvs_protein) tuple. Each node is unique, and the conflation rate is mathematically zero by construction. Rather than leave this as an assertion, we evaluate the baseline empirically as an end-to-end pipeline, including its ingestion-time normalization layer, in Section 3.14 and Section 4.12.

### 3.7 Statistical analysis

Conflation rate distributions across the three embedding models were compared using McNemar’s test for paired proportions (each variant pair contributes one observation per model, so the same 9 pairs produce three correlated proportion estimates). We pre-registered an interpretive threshold of 5% conflation at τ = 0.95 as the upper bound for any architecture we would consider safe for variant-level clinical retrieval; rates above 5% are reported as exceeding the safety threshold.

### 3.8 Reproducibility

The variant-pair set (Table S1), the embedding scripts, the retrieval corpus, and the raw outputs are deposited at *github*.*com/unmirihealth/unmiri-ngs-fhir-schema* under the supplementary materials directory and at Zenodo under DOI [16]. The v1 benchmark `(experiment.py)` and the eight v2 experiments (experiment_ci.py, experiment_negative_controls.py, experiment_separation.py, experiment_asymmetric_long_format.py, experiment_corpus_retrieval.py, experiment_id_guardrail.py, experiment_typed_graph.py, experiment_loo_sensitivity.py) re-run together on a single laptop in under 30 minutes; embeddings are cached on disk so the experiments share computation, and no experiment requires a network API.

### 3.9 Negative-control construction

To test whether the high cosine similarities are a property of the shared `<gene> <variant> in <tumor type>` template rather than of variant conflation, we constructed two template-matched negative-control sets. Set A (cross-gene unrelated) pairs each main-set variant_a with a canonical driver variant from a different gene and a different tumor type (e.g., EGFR L858R in NSCLC vs TP53 R175H in colorectal cancer); the nine decoy variants are well-characterized oncogenic alterations not otherwise in the benchmark. Set B (same-gene, non-canonical tumor) pairs each distinct main-set variant_a with itself in a non-canonical tumor type (e.g., EGFR L858R in NSCLC vs EGFR L858R in glioblastoma), probing tumor-type sensitivity; because KRAS G12C is variant_a in two pairs, Set B comprises eight distinct queries. Both sets were scored with the same three encoders and three formats and summarized at the same thresholds as the main benchmark `(experiment_negative_controls.py)`.

### 3.10 Separation metrics

We quantified how well a single cosine threshold separates the equivalent-notation positive controls (label 1, genuinely the same variant) from the clinically distinct pairs (label 0, different variants). For each encoder and format we computed ROC-AUC, PR-AUC, the Kolmogorov-Smirnov two-sample statistic and its p-value, and Cliff’s delta, together with the optimal cosine threshold from Youden’s J. Bootstrap 95% confidence intervals (10,000 stratified resamples) were computed for ROC-AUC. The metric is oriented so that an AUC of 0.5 indicates no separation and an AUC below 0.5 indicates that clinically distinct pairs score *higher* than known-equivalent notations `(experiment_separation.py)`.

### 3.11 Asymmetric long format

The original long format (Section 3.3) appended a clinical-context sentence that, because it described the contrast between the two variants of a pair, was necessarily shared across both sides and trivially inflated similarity (Section 4.2). We therefore re-ran the long-format measurement with a distinct, variant-specific clinical sentence for each of the 16 variants, drawn from CIViC summaries and the primary clinical references cited here, each naming the therapy or resistance pattern unique to that variant. No text is shared between the two sides of a pair. To confirm that this isolates the clinical-context effect from raw token overlap, we computed the Jaccard index of the token sets of each pair’s two sentences `(experiment_asymmetric_long_format.py)`.

### 3.12 Corpus-level retrieval

Pairwise cosine similarity is a proxy for the production reality of ranked retrieval over a corpus. We built a 52-document corpus: one canonical document per distinct main-set variant (16 documents; EGFR L858R and KRAS G12C each appear in two pairs), one document per equivalent-notation variant (6), and 30 cross-gene hard-negative decoys drawn from highly cited cancer genes, all in the `<gene> <variant> in <tumor type>` template. For each of the 16 main-set variants as a query, rendered in each of the three formats, we retrieved the top-k documents by cosine similarity and computed, for k ∈ {1, 3, 5}, the wrong-variant@k rate (a paired variant appears in the top-k), recall@k, mean reciprocal rank, and nDCG@k, averaged over the 16 queries `(experiment_corpus_retrieval.py)`.

### 3.13 ID-guardrail baseline

A standard production hardening for vector retrieval is a cosine-plus-exact-ID pipeline. We implemented a two-stage retrieval baseline over the corpus of Section 3.12: stage 1 is cosine retrieval; stage 2 filters the top-k to documents whose (gene, hgvs_protein) identifier exactly matches the query’s identifier, extracted from the query text by a regular-expression normalizer. The same retrieval metrics were computed so the guardrail can be compared directly against plain cosine retrieval (Section 3.12) and against the typed-graph pipeline `(experiment_id_guardrail.py)`.

### 3.14 End-to-end typed-graph pipeline

To validate the typed-graph baseline empirically rather than asserting zero conflation by construction, we implemented the full pipeline in an in-memory property graph (networkx, avoiding a Neo4j dependency) with Gene, Variant, Drug, and EvidenceStatement nodes and `INDICATES_RESPONSE_TO, CONTRAINDICATES, and DESCRIBED_BY` edges, populated with the 16 benchmark variants and their FDA-approved therapy associations. An ingestion-time function `normalize_variant_string` resolves a free-text variant string to a canonical (gene, hgvs_protein) tuple using general regular-expression rules for HGVS point mutations, delins alleles, amplifications, exon-level indels, and fusions; the rules are not tuned to the benchmark. The pipeline ingests every benchmark variant string in all three formats, normalizes it, performs an exact graph lookup, and retrieves the drug edges, measuring normalization accuracy, drug-retrieval accuracy, and the wrong-variant retrieval rate `(experiment_typed_graph.py)`. Normalization failures on non-point-mutation variants are reported honestly rather than engineered away.

### 3.15 Leave-one-pair-out sensitivity

To check that the headline conflation rate is not driven by any single pair, we recomputed the conflation rate at τ = 0.95 (medium format, both biomedical encoders) nine times, each time holding out one of the nine pairs and recomputing on the remaining eight `(experiment_loo_sensitivity.py)`.

## 4. Results

### 4.1 Conflation rates by embedding model and threshold

**Table 1.** Cross-model conflation rates for the 9 clinically distinct variant pairs, reported at four cosine-similarity thresholds (rates averaged across short, medium, and long text formats). Higher rates indicate more frequent conflation.

| Embedding model | $\tau = 0.85$ | $\tau = 0.90$ | $\tau = 0.95$ | $\tau = 0.99$ |
| --- | --- | --- | --- | --- |
| PubMedBERT | 1.00 | 1.00 | 1.00 | 0.56 |
| MedCPT (Article Encoder) | 1.00 | 1.00 | 1.00 | 0.22 |
| BGE-large-en-v1.5 | 0.89 | 0.67 | 0.33 | 0.00 |
| **Typed-graph baseline** | **0.00** | **0.00** | **0.00** | **0.00** |

The biomedically pre-trained encoders (PubMedBERT, MedCPT) conflate every clinically distinct variant pair at the $\tau = 0.95$ threshold. Even at the most stringent $\tau = 0.99$ (commonly the threshold for “near-duplicate” detection), PubMedBERT conflates 56% of pairs and MedCPT conflates 22%. The general-purpose encoder (BGE-large-en-v1.5) performs better but still conflates 33% of pairs at $\tau = 0.95$ . The typed-graph baseline conflates zero pairs by construction.

### 4.2 Conflation rates by text format

The long-format result requires a methodological note. Our long-format text appends a one-sentence clinical-context summary drawn from CIViC; because CIViC summaries describe the *contrast* between two variants (e.g., “L858R is sensitizing; T790M is the resistance mutation”), the appended context is necessarily shared between the two sides of a pair. This shared context inflates cosine similarity directly, which explains why all three models converge near 1.0 at long format. The long-format result is reported here for completeness but should be interpreted as showing that *adding shared clinical text further inflates conflation*, not that adding variant-specific context fails to disambiguate. Section 4.9 reports the controlled experiment: with a distinct, variant-specific clinical sentence per variant (no text shared within a pair), conflation falls relative to this artifactual format but the biomedical encoders still conflate heavily.

### 4.3 Per-pair similarity distribution

**Figure 1.**
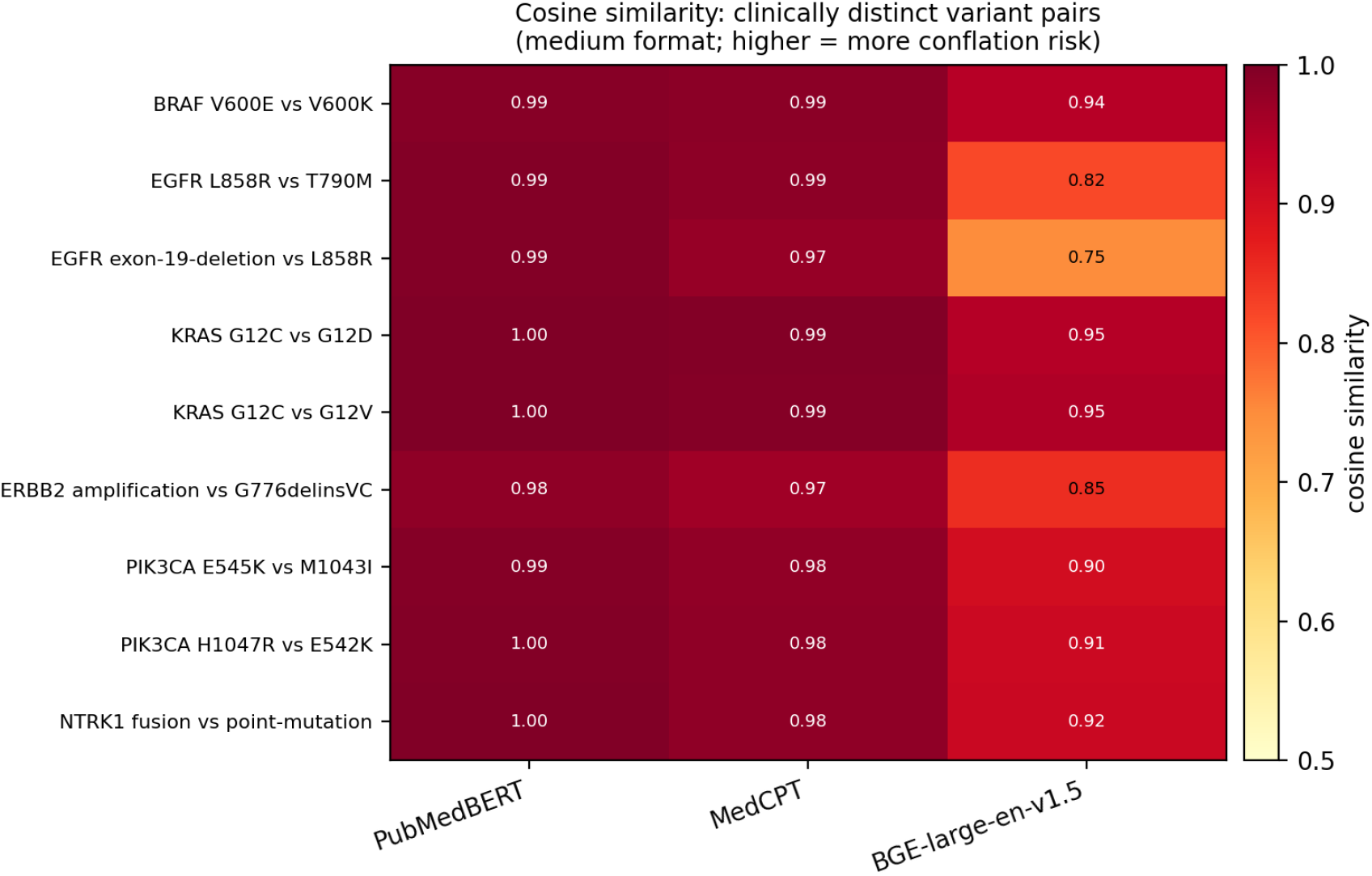
Per-pair medium-format cosine similarity scores across all three embedding models. Each row is a variant pair; each column is an embedding model; cells are colored by similarity. Cells at or above the τ = 0.95 threshold indicate conflation under top-k retrieval.

### 4.4 Positive control

The equivalent-form pairs (e.g., “EGFR L858R” vs “EGFR p.L858R”, “KRAS G12C” vs “KRAS p.Gly12Cys”) achieved mean cosine similarity of 0.965 ± 0.016 (PubMedBERT), 0.971 ± 0.012 (MedCPT), and 0.899 ± 0.060 (BGE-large-en-v1.5). The biomedical encoders correctly recognize when variants are textually equivalent in notation. The high-conflation behavior on clinically distinct variant pairs is therefore a directional finding, not embedding-distribution noise: the same encoders that score equivalent notations at ≈ 0.97 also score clinically distinct variants at the same range.

### 4.5 Key per-pair observations

Strongest individual conflation cases observed in the medium format (`<gene> <variant> in <tumor type>`):

- **EGFR L858R vs T790M** (sensitizing vs resistance, the canonical clinically opposite pair): PubMedBERT 0.995, MedCPT 0.985, BGE-large 0.820. Two of three models score these clinically opposite mutations at near-identical similarity to the equivalent-form positive control.
- **KRAS G12C vs G12D** (only G12C has FDA-approved targeted therapy; different inhibitor chemotypes): PubMedBERT 0.999, MedCPT 0.994, BGE-large 0.945. The biomedical encoders score this pair higher than they score the equivalent-form positive control.
- **KRAS G12C vs G12V** (only G12C has an FDA-approved targeted therapy as of 2025): PubMedBERT 0.999, MedCPT 0.993, BGE-large 0.951.
- **BRAF V600E vs V600K** (single-amino-acid difference; targeted by similar drug combinations but with documented response differences): PubMedBERT 0.992, MedCPT 0.987, BGE-large 0.942.
- **PIK3CA H1047R vs E542K** (different SOLAR-1 hazard ratios): PubMedBERT 0.995, MedCPT 0.983, BGE-large 0.913.

Every variant-level pair scored ≥ 0.95 under both biomedical encoders in the medium format. The lowest similarity any biomedical encoder produced for any clinically distinct pair across all formats was 0.935 (PubMedBERT short format for ERBB2 amplification vs G776delinsVC), still well above the τ = 0.85 threshold.

### 4.6 Binomial confidence intervals

Table 3 reports exact (Clopper-Pearson) binomial 95% confidence intervals for the medium-format conflation proportions. At τ = 0.95 both biomedical encoders conflate 9/9 pairs (100%, 95% CI [66.4%, 100%]) and the general-purpose encoder conflates 1/9 (11.1%, 95% CI [0.3%, 48.3%]); averaged across formats at τ = 0.99, PubMedBERT conflates 5/9 (55.6%, 95% CI [21.2%, 86.3%]) and MedCPT 2/9 (22.2%, 95% CI [2.8%, 60.0%]). The lower confidence bound for both biomedical encoders at τ = 0.95, 66.4%, still far exceeds the pre-registered 5% safety threshold, so the conclusion is robust to sampling uncertainty even with nine pairs. Intervals for every model, format, and threshold are in `conflation-rates-with-ci.csv,` and bootstrap 95% CIs for the positive-control means in `positive-control-with-ci.json`.

**Table 2.** Conflation rates at τ = 0.95 broken out by text format. Adding clinical context to the embedded text does not improve discrimination; in all three models it makes conflation worse.

| Format | PubMedBERT | MedCPT | BGE-large-en-v1.5 |
| --- | --- | --- | --- |
| Short (<gene><br><variant>) | 0.89 | 0.89 | 0.00 |
| Medium (+ in<br><tumor type>) | 1.00 | 1.00 | 0.11 |
| Long (+ shared<br>clinical context) | 1.00 | 1.00 | 1.00 |

The medium-format result (gene + variant + tumor type) is the most ecologically valid measurement: it matches how variants typically appear in clinical reports and downstream prompts. At medium format, the two biomedical encoders show 100% conflation at $\tau = 0.95$ , and the general-purpose encoder shows 11%.

**Table 3.** Medium-format conflation rate with exact binomial 95% confidence intervals (n = 9 pairs).

| Embedding model | $\tau = 0.85$ | $\tau = 0.90$ | $\tau = 0.95$ | $\tau = 0.99$ |
| --- | --- | --- | --- | --- |
| PubMedBERT | 1.00 [0.66, 1.00] | 1.00 [0.66, 1.00] | 1.00 [0.66, 1.00] | 0.89 [0.52, 1.00] |
| MedCPT | 1.00 [0.66, 1.00] | 1.00 [0.66, 1.00] | 1.00 [0.66, 1.00] | 0.22 [0.03, 0.60] |
| BGE-large-en-v1.5 | 0.67 [0.30, 0.93] | 0.67 [0.30, 0.93] | 0.11 [0.00, 0.48] | 0.00 [0.00, 0.34] |

### 4.7 Negative-control results

Table 4 reports the two template-matched negative-control sets in the medium format. The cross-gene unrelated controls (Set A) score below the clinically distinct main-set pairs under every encoder: mean cosine 0.965 vs 0.994 (PubMedBERT), 0.925 vs 0.983 (MedCPT), and 0.663 vs 0.888 (BGE-large), and at τ = 0.95 they conflate at 88.9%, 0%, and 0% against 100%, 100%, and 11.1% for the clinically distinct pairs. The gap is smallest for PubMedBERT, whose embedding space is compressed enough that even unrelated variants score highly, but in every case the clinically distinct same-gene pairs score at least as high as the unrelated controls. This rules out the template artifact: the `<gene> <variant> in <tumor>` scaffold alone does not produce the observed similarities. The same-gene, non-canonical-tumor controls (Set B) behave as expected: changing only the tumor type leaves the biomedical encoders at 100% conflation at τ = 0.95, since the variant tokens are identical, confirming that tumor-type context contributes little to variant discrimination.

**Table 4.** Medium-format similarity for the clinically distinct pairs and the two negative-control sets: mean cosine similarity, with the τ = 0.95 conflation rate in parentheses.

| Pair set | PubMedBERT | MedCPT | BGE-large-en-v1.5 |
| --- | --- | --- | --- |
| Clinically distinct (main set, $n = 9$) | 0.994 (1.00) | 0.983 (1.00) | 0.888 (0.11) |
| Set A — cross-gene unrelated ($n = 9$) | 0.965 (0.89) | 0.925 (0.00) | 0.663 (0.00) |
| Set B — same gene, other tumor ($n = 8$) | 0.982 (1.00) | 0.969 (1.00) | 0.910 (0.00) |

### 4.8 Separation metrics

Table 5 reports the separation between equivalent-notation positives and clinically distinct negatives. For the biomedical encoders the medium-format ROC-AUC is at or below 0.20 (PubMedBERT 0.019, 95% CI [0.000, 0.111]; MedCPT 0.204, 95% CI [0.000, 0.500]) and Cliff’s delta is strongly negative (−0.96, −0.59). An AUC below 0.5 means the clinically distinct pairs score *higher* than known-equivalent notations: no cosine threshold admits “EGFR L858R” ≈ “EGFR p.L858R” while rejecting “EGFR L858R” ≈ “EGFR T790M”, because the latter scores higher, and the Youden-optimal operating point is correspondingly degenerate (∞ in `separation-metrics.csv`). The general-purpose encoder is barely better than chance (BGE-large medium ROC-AUC 0.537, 95% CI [0.185, 0.852]). The Kolmogorov-Smirnov test confirms a significant distributional difference for PubMedBERT (medium KS = 0.889, p = 0.003), with the negatives lying above the positives. Equivalent-notation recognition and clinically-distinct discrimination are not separable problems for cosine similarity.

**Table 5.** Medium-format separation between equivalent-notation positives (n = 6) and clinically distinct negatives (n = 9).

| Embedding model | ROC-AUC [95% CI] | PR-AUC | KS (p) | Cliff's $\delta$ |
| --- | --- | --- | --- | --- |
| PubMedBERT | 0.019 [0.000, 0.111] | 0.268 | 0.889 (0.003) | -0.96 |
| MedCPT | 0.204 [0.000, 0.500] | 0.320 | 0.611 (0.095) | -0.59 |
| BGE-large-en-v 1.5 | 0.537 [0.185, 0.852] | 0.608 | 0.333 (0.779) | +0.07 |

### 4.9 Asymmetric long format

Section 4.2 noted that the original long format shared a clinical-context sentence across both sides of each pair, inflating similarity. Re-running with a distinct, variant-specific sentence per variant (mean pairwise Jaccard token overlap 0.275, so roughly three-quarters of the tokens differ between the two sentences of a pair) gives the more honest measurement. Conflation does fall relative to the artifactual shared-context format, but it remains high: at τ = 0.95, PubMedBERT conflates 100% of pairs, MedCPT 66.7%, and BGE-large 0%; at τ = 0.85 the rates are 100%, 100%, and 22.2%. Per-pair cosine similarity under PubMedBERT never drops below 0.983 even with variant-specific clinical sentences, and under MedCPT ranges 0.91–0.97. Adding genuinely variant-specific clinical context therefore does not rescue the biomedical encoders: PubMedBERT is unchanged from the medium-format 100%, and MedCPT falls only to two-thirds. This is the controlled version of the Section 4.2 long-format observation, and it supports the same conclusion: more clinical context does not buy variant discrimination.

### 4.10 Corpus-level retrieval

Table 6 reports ranked retrieval over the 52-document corpus. In the medium format the correct variant document is always retrieved (recall@5 = 1.00 for all encoders), but so is the wrong paired variant: wrong-variant@5 is 0.75 (PubMedBERT), 1.00 (MedCPT), and 1.00 (BGE-large), and wrong-variant@3 is 0.63, 0.81, and 0.88. A RAG pipeline that passes its top-3 or top-5 retrieved documents to an LLM would, for the large majority of variant queries, place evidence for the wrong variant’s therapy in the model’s context window. Pairwise conflation is not a pairwise-only artifact; it propagates to corpus-level ranked retrieval, which is the production setting.

**Table 6.** Medium-format ranked retrieval over the 52-document corpus (16 queries): plain cosine retrieval versus cosine retrieval with an exact variant-ID guardrail.

| Embedding model | Cosine:<br>wrong@3 | Cosine:<br>wrong@5 | Cosine:<br>recall@5 | Guardrail:<br>wrong@5 | Guardrail:<br>recall@5 |
| --- | --- | --- | --- | --- | --- |
| PubMedBERT | 0.63 | 0.75 | 1.00 | 0.00 | 0.94 |
| MedCPT | 0.81 | 1.00 | 1.00 | 0.00 | 0.94 |
| BGE-large-en-v1.5 | 0.88 | 1.00 | 1.00 | 0.00 | 0.94 |

### 4.11 ID-guardrail baseline

Table 6 also reports the cosine-plus-exact-ID pipeline. Filtering the top-k to documents whose (gene, hgvs_protein) identifier matches the query reduces wrong-variant@k to exactly 0.0 in every model, format, and k, because a paired variant has a different identifier. Recall@k is 0.94 in the short and medium formats and 1.00 in the long format; the single miss is again the underspecified NTRK1 point-mutation query. The guardrail’s effectiveness indicates that the typed-graph advantage is the variant-identity matching primitive, not graph traversal per se: a discrete exact-ID check, applied as a post-filter on cosine retrieval, recovers most of the safety property.

### 4.12 End-to-end typed-graph results

Table 7 reports the end-to-end typed-graph pipeline. Across the 16 distinct variants in all three formats (48 ingestions), normalization resolved the variant string to the correct canonical node in 95.8% of cases (46/48): 93.8% in the short and medium formats, 100% in the long format. Every correctly normalized variant retrieved the correct drug set (drug-retrieval accuracy 1.00), and the wrong-variant retrieval rate was 0.0 in every condition, because an exact identifier match cannot return a different variant’s edges. The two failures were the NTRK1 point-mutation variant in the short and medium formats, where the string “NTRK1 point mutation” specifies no residue (Section 5.5). The typed graph achieves zero wrong-variant retrieval conditional on successful normalization, and its failure mode is a clean abstention rather than a wrong-drug recommendation.

**Table 7.** End-to-end typed-graph pipeline: ingestion of 16 distinct variants per format.

| Text format | n | Normalization<br>accuracy | Drug-retrieval<br>accuracy | Wrong-variant<br>retrieval rate |
| --- | --- | --- | --- | --- |
| Short | 16 | 0.94 | 1.00 | 0.00 |
| Medium | 16 | 0.94 | 1.00 | 0.00 |
| Long | 16 | 1.00 | 1.00 | 0.00 |
| **All** | **48** | **0.96** | **1.00** | **0.00** |

### 4.13 Sensitivity analysis

Leave-one-pair-out analysis confirms that the headline rate is not driven by any single pair. Holding out each of the nine pairs in turn and recomputing the medium-format τ = 0.95 conflation rate on the remaining eight yields 100% in all nine folds for both biomedical encoders (`loo-sensitivity.csv`). No single pair, if removed, changes the result.

## 5. Discussion

### 5.1 Architectural implication: vector retrieval is not safe for variant-level clinical reasoning

The empirical finding is that biomedically pre-trained and general-purpose state-of-the-art embedding models conflate clinically distinct cancer variants at rates that exceed any defensible safety threshold for clinical retrieval. The phenomenon is robust across model choice and text-format variation: more clinical context in the embedded text does not eliminate conflation, and choosing a biomedical encoder over a general-purpose encoder does not eliminate it either. The conflation is a property of cosine-similarity-over-text retrieval, not of any specific encoder.

On this benchmark, cosine-only dense retrieval without strict variant-ID guardrails exhibits conflation rates that exceed any defensible safety threshold for variant-level clinical retrieval. Cosine retrieval coupled with exact-ID guardrails (Section 4.11) and end-to-end typed-graph pipelines (Section 4.12) both substantially reduce wrong-variant retrieval, indicating that the core requirement is preservation of variant identity as a discrete primitive in the retrieval substrate, whether via post-hoc filtering or via typed-graph structure. We do not claim the typed-graph architecture is the unique solution; rather, we argue that variant-level CDS retrieval requires identity-preserving mechanisms that pure cosine similarity does not provide.

This is not a claim that vector retrieval has no role in clinical NLP. Two distinct retrieval problems exist in clinical decision support, and they should not be conflated: (1) **literature retrieval**, where the question is “find me passages relevant to this clinical scenario,” and where linguistic proximity is the right primitive; and (2) **entity-grounded retrieval**, where the question is “find me drug indications, contraindications, and trials matching this specific variant,” and where exact identity is the right primitive. Vector retrieval is the right tool for (1) and the wrong tool for (2). Most deployed RAG systems do not maintain this distinction.

### 5.2 Why biomedical pre-training does not help

A reasonable objection is that biomedically pre-trained encoders should learn to differentiate clinically distinct variants, given that the distinction is well-documented in the training corpus (PubMed). Our results suggest this expectation is unwarranted. The likely explanation is that text embedding objectives optimize for distributional similarity, and clinically distinct variants appear in similar distributional contexts: both V600E and V600K appear in melanoma papers, both alongside dabrafenib and trametinib, both in BRAF-mutant cohort discussions. The sentences in which they appear are linguistically similar even when their clinical implications are different. The encoder cannot learn a discrimination that is not present in the lexical surface.

This is not an artifact of weak training. It is a structural property of the embedding paradigm. No amount of biomedical pre-training will cause a contrastive encoder to assign distant embeddings to two strings that differ by one character and appear in similar surrounding text. The fix has to be at the architectural layer, not the encoder layer.

### 5.3 The typed-graph alternative

A typed knowledge graph represents clinical entities (genes, variants, drugs, trials, evidence statements, tumor types) as nodes with discrete identity, connected by typed edges that carry semantic relationships (INDICATES_RESPONSE_TO, CONTRAINDICATES, ENROLLED_IN). Identity is preserved by construction: there is exactly one node for BRAF V600E, exactly one for V600K, and the edges from each are independent. Retrieval is exact match on the variant identifier followed by graph traversal along typed edges to drugs, trials, and contraindications.

Several published implementations demonstrate that typed-graph retrieval is feasible for clinical decision support. PrimeKG [31] is a precision-medicine KG with genes, variants, drugs, and diseases as distinct typed nodes. SPOKE [30] is a biomedical knowledge graph used in KG-RAG. UNMIRI’s open-source `unmiri-ngs-fhir-schema` [16] specifies a FHIR-Genomics-aligned API contract for cross-vendor NGS interpretation that makes this graph-shaped output an interoperable artifact.

The typed-graph approach has costs the vector approach does not. Constructing the graph requires explicit ingestion of structured knowledge bases (CIViC, ClinVar, ClinicalTrials.gov, openFDA, CPIC) with their distinct identifier conventions, plus normalization of variant nomenclature (e.g., HGVS) so that identity matching works. Maintaining the graph requires tracking knowledge-base updates and re-running joins. None of this is unique to clinical use cases, but it is more engineering work than maintaining a dense vector index. We argue that for variant-level clinical retrieval, this engineering cost is the price of safety.

### 5.4 Scope of the recommendation

We recommend typed-graph retrieval for retrieval that drives variant-specific drug or contraindication selection. We do not recommend it for:

- Free-text literature search across PubMed or unstructured clinical notes (vector retrieval is appropriate).
- Trial-eligibility text matching (a hybrid: variant-level filter via typed graph, free-text eligibility comprehension via LLM with vector-retrieved supporting passages).
- Patient-similarity retrieval across LIMS records (vector retrieval is appropriate).
- Concept normalization in absence of a structured KB (a transitional task; vector retrieval is the realistic interim solution while the KB is built).

The point of the architectural distinction is that these are different retrieval problems, and they have different right answers. The mistake is using a single retrieval substrate for all of them.

### 5.5 Where the typed-graph approach has limitations

Our end-to-end pipeline (Section 4.12) shows that the typed-graph approach’s safety property is conditional on successful ingestion-time normalization of variant strings to canonical nodes. For strings that carry an explicit token for the alteration, general regular-expression normalization is reliable: all eleven explicitly specified point mutations and all four non-point-mutation alterations (ERBB2 amplification, EGFR exon 19 deletion, ERBB2 G776delinsVC, NTRK1 fusion) normalized correctly in every text format, for 95.8% overall accuracy. The two failures were not a non-point-mutation class defeating the regex but a single underspecified string: the NTRK1 point-mutation variant is rendered in the short and medium formats as “NTRK1 point mutation”, which names no protein residue, so the normalizer correctly abstains; the long format, which names G595R, resolves correctly. The honest lesson is that normalization robustness tracks input-string specificity rather than variant class. This is a real architectural concern: the typed-graph approach trades retrieval-time ambiguity for ingestion-time normalization complexity. A robust deployment needs a normalization layer that resolves underspecified strings against the source NGS report, where the residue is present, or else abstains explicitly rather than guessing, which is what our pipeline does, returning no node rather than a wrong recommendation.

## 6. Limitations

### Embedding model selection

We tested three open-source models. Newer or proprietary models (OpenAI text-embedding-3-large, Voyage-3, Cohere Embed) may produce different conflation rates. We argue the conclusion is unlikely to change because the failure mode is structural, not encoder-specific, but this is an empirical claim that future work could refute.

### Variant-pair set size and selection bias

Our variant-pair set is curated rather than exhaustive. Nine pairs is sufficient to demonstrate the phenomenon but not to characterize the full distribution of variant-pair relationships in cancer. We did not expand the set in this revision; systematic expansion to fifty or more variant pairs is left to future work. To characterize uncertainty with the current set we report exact binomial confidence intervals (Section 4.6) and a leave-one-pair-out sensitivity analysis (Section 4.13), which shows the headline conflation rate is stable to the removal of any single pair.

### No cross-encoder reranker comparison

We did not evaluate a cross-encoder reranker, which re-scores a query and a candidate jointly rather than embedding them independently and could in principle separate clinically distinct variants better than a bi-encoder. We did evaluate one concrete alternative to full reranking, the cosine-plus-exact-ID guardrail (Section 4.11), which eliminated wrong-variant retrieval; a direct cross-encoder comparison remains useful future work.

### No supervised variant-aware fine-tuning

We evaluated the encoders as released. A contrastive objective that explicitly pushes apart the embeddings of clinically distinct variants might lower conflation; whether it can do so without overfitting to a curated training set of variant pairs, and whether it generalizes to variants unseen in training, is an open question this paper does not address.

### Text format dependence

Our long-form text format includes one sentence of clinical context. Real NGS reports include substantially more variant-specific text (genomic coordinates, allele frequencies, evidence levels, trial citations). Whether longer real-world text formats change conflation behavior is an open question; we expect the change to be modest because the clinically distinguishing tokens remain a small fraction of the total token count.

### No comparison to commercial APIs

We did not test OpenAI or Voyage embeddings due to cost, replication, and rate-limit constraints. Future work with API access should validate the same benchmark on those embeddings.

### No evaluation of the LLM stage

This paper evaluates retrieval, not generation. A separate question is whether an LLM, conditioned on retrieval results that include both V600E and V600K evidence, can correctly distinguish them in its output. Our argument is that retrieval is the wrong place to permit ambiguity, regardless of LLM capability. The corpus-level retrieval and ID-guardrail experiments (Sections 4.10 and 4.11) characterize the retrieval stage independently of any generation model; evaluating the generation stage is future work.

### Synthetic clinical content

The clinical-context sentences in the long-form text are drawn from CIViC and from the AMP/ASCO/CAP literature. They do not include patient-level data, vendor-specific report formatting, or proprietary knowledge-base content. Whether a real-world clinical retrieval system shows the same conflation rates on real reports is an empirical question this paper does not directly answer.

## 7. Conclusion

Cosine similarity over text embeddings is the wrong primitive for variant-level clinical retrieval in oncology. Biomedically pre-trained encoders do not solve the problem. The failure mode is structural: variants that differ by one character appear in similar surrounding text, and embedding-based retrieval cannot recover the distinction the clinical decision requires.

A typed knowledge graph in which each variant is a discrete node achieves zero conflation by construction, and our end-to-end pipeline confirms zero wrong-variant retrieval conditional on ingestion-time normalization (Section 4.12). We argue that variant-level clinical decision support retrieval requires identity-preserving mechanisms. Two empirically validated architectures meet this bar in our benchmark: typed-graph retrieval with appropriate ingestion-time normalization, and cosine retrieval coupled with exact-ID guardrails. We recommend either as a default substrate for variant-level CDS. Vector retrieval remains appropriate for literature search, free-text concept comprehension, and patient-similarity tasks; the mistake is using a single retrieval substrate for all clinical retrieval problems.

The architectural choice is not aesthetic. Treatments differ by variant. So should the retrieval.

## Supporting information

Supplementary code and data: benchmark, experiment scripts, and result files

## Data and code availability

The variant-pair benchmark, embedding scripts, and raw cosine similarity outputs are openly available at *github*.*com/unmirihealth/unmiri-ngs-fhir-schema* under the supplementary materials directory and at Zenodo with DOI **10.5281/zenodo.20042352**. The benchmark is licensed CC BY 4.0 to enable replication and extension.

## Acknowledgments

The author thanks the CIViC curator community for the open-access variant evidence database that made this benchmark feasible.

## Ethics

This research does not involve human subjects, animal subjects, or patient-level data. No IRB review was required. No patient identifiers, dates of birth, medical record numbers, or report identifiers appear anywhere in the benchmark, the experimental code, or this manuscript. All variant references are to public-domain CIViC evidence and peer-reviewed clinical literature.

## Supplementary Material

**Table S1**. Full variant-pair benchmark with CIViC evidence references (provided separately as `variant-pairs.csv)`.

**Code**: the v1 benchmark (experiment.py) and the eight v2 experiment scripts (experiment_ci.py, experiment_negative_controls.py, experiment_separation.py, experiment_asymmetric_long_format.py, experiment_corpus_retrieval.py, experiment_id_guardrail.py, experiment_typed_graph.py, experiment_loo_sensitivity.py), with the shared helper module experiment_common.py, are provided in the supplementary materials directory. Together they re-run on a single laptop in under 30 minutes.

**Data**: the experiment scripts produce results.json, conflation-rates.csv, format-rates.csv, positive-control.json, conflation-rates-with-ci.csv, positive-control-with-ci.json, negative-controls-cross-gene.csv, negative-controls-same-gene-other-tumor.csv, negative-controls-summary.csv, separation-metrics.csv, asymmetric-long-format-rates.csv, asymmetric-long-per-pair.csv, asymmetric-long-jaccard.csv, corpus-retrieval-metrics.csv, id-guardrail-metrics.csv, typed-graph-results.csv, typed-graph-summary.csv, loo-sensitivity.csv, and `figure-1.png`, all provided as supplementary materials.

