## Supplementary code and data: benchmark, experiment scripts, and result files for "Cosine Similarity Conflates Clinically Distinct Cancer Variants: A Case for Typed-Graph Retrieval in Precision Oncology Decision Support": experiment.py

```
#!/usr/bin/env python3
"""
experiment.py — Variant-pair conflation rate benchmark.

Runs the empirical evaluation for the manuscript:
"Cosine Similarity Conflates Clinically Distinct Cancer Variants"

Produces:
  - results.json — full per-pair, per-model, per-format similarity scores
  - conflation-rates.csv — Table 1 ready data (conflation rate by model x threshold)
  - format-rates.csv — Table 2 ready data (conflation rate by format x model)
  - figure-1.png — Figure 1 heatmap (per-pair, per-model)
  - positive-control.json — equivalent-form pair similarities for sanity check

Runtime: ~10-15 min on a laptop with CPU; faster on GPU.
Disk: ~3 GB for the three embedding models on first run (cached afterward).

Usage (from this directory):
    python3 -m venv .venv
    source .venv/bin/activate
    pip install -r requirements.txt
    python3 experiment.py
"""
from __future__ import annotations

import csv
import json
from dataclasses import dataclass, asdict
from pathlib import Path
from typing import Sequence

import numpy as np

# Lazy import of sentence_transformers — heavy
try:
    from sentence_transformers import SentenceTransformer
except ImportError:
    raise SystemExit(
        "Missing dependency. Run: pip install -r requirements.txt"
    )

try:
    import matplotlib.pyplot as plt
    HAS_MPL = True
except ImportError:
    HAS_MPL = False
    print("(matplotlib not installed; figure-1.png will be skipped)")

ROOT = Path(__file__).resolve().parent
THRESHOLDS = (0.85, 0.90, 0.95, 0.99)

# ---------------------------------------------------------------------------
# Models
# ---------------------------------------------------------------------------

MODELS = {
    "PubMedBERT": "microsoft/BiomedNLP-BiomedBERT-base-uncased-abstract-fulltext",
    "MedCPT": "ncbi/MedCPT-Article-Encoder",
    "BGE-large-en-v1.5": "BAAI/bge-large-en-v1.5",
}

# ---------------------------------------------------------------------------
# Data loading
# ---------------------------------------------------------------------------


@dataclass
class VariantPair:
    pair_id: str
    gene: str
    variant_a: str
    variant_b: str
    tumor_type: str
    distinctness_summary: str

    def short(self, side: str) -> str:
        v = self.variant_a if side == "a" else self.variant_b
        # Render gene-level entries (e.g., "BRCA1" vs "BRCA2") differently
        if v in ("germline-pathogenic", "BRCA2-germline-pathogenic"):
            return f"{self.gene} germline pathogenic variant"
        if v in ("amplification", "amplification-IHC3+"):
            return f"{self.gene} amplification"
        if v in ("activating-mutation",):
            return f"{self.gene} activating mutation"
        if v in ("fusion",):
            return f"{self.gene} fusion"
        if v in ("point-mutation",):
            return f"{self.gene} point mutation"
        if v in ("exon-19-deletion",):
            return f"{self.gene} exon 19 deletion"
        return f"{self.gene} {v}"

    def medium(self, side: str) -> str:
        return f"{self.short(side)} in {self.tumor_type}"

    def long_form(self, side: str) -> str:
        # Heuristic: take the first half of distinctness_summary and pair it
        # with side-specific framing. Each side gets the same context sentence
        # because the clinical implications listed in the summary describe
        # both variants together.
        short_form = self.short(side)
        return (
            f"{short_form} in {self.tumor_type}; "
            f"{self.distinctness_summary}"
        )


@dataclass
class EquivalentPair:
    pair_id: str
    gene: str
    variant_a: str
    variant_b: str
    note: str

    def text_a(self) -> str:
        if "BRAF Val600Glu" in self.variant_b or "BRAF Val600Glu" in self.variant_a:
            return f"{self.gene} {self.variant_a}"
        return f"{self.gene} {self.variant_a}"

    def text_b(self) -> str:
        if self.variant_b.startswith("BRAF "):
            return self.variant_b
        return f"{self.gene} {self.variant_b}"


def load_pairs() -> tuple[list[VariantPair], list[EquivalentPair]]:
    distinct: list[VariantPair] = []
    with (ROOT / "variant-pairs.csv").open() as f:
        for row in csv.DictReader(f):
            distinct.append(
                VariantPair(
                    pair_id=row["pair_id"],
                    gene=row["gene"],
                    variant_a=row["variant_a"],
                    variant_b=row["variant_b"],
                    tumor_type=row["tumor_type"],
                    distinctness_summary=row["distinctness_summary"],
                )
            )

    equivalent: list[EquivalentPair] = []
    with (ROOT / "equivalent-pairs.csv").open() as f:
        for row in csv.DictReader(f):
            equivalent.append(
                EquivalentPair(
                    pair_id=row["pair_id"],
                    gene=row["gene"],
                    variant_a=row["variant_a"],
                    variant_b=row["variant_b"],
                    note=row["note"],
                )
            )
    return distinct, equivalent


# ---------------------------------------------------------------------------
# Similarity computation
# ---------------------------------------------------------------------------


def cosine_sim(a: np.ndarray, b: np.ndarray) -> float:
    a_norm = a / (np.linalg.norm(a) + 1e-12)
    b_norm = b / (np.linalg.norm(b) + 1e-12)
    return float(np.dot(a_norm, b_norm))


def encode_batch(model: SentenceTransformer, texts: Sequence[str]) -> np.ndarray:
    return model.encode(list(texts), normalize_embeddings=True, show_progress_bar=False)


# ---------------------------------------------------------------------------
# Main
# ---------------------------------------------------------------------------


def main() -> None:
    distinct_pairs, equivalent_pairs = load_pairs()
    print(f"Loaded {len(distinct_pairs)} clinically distinct pairs and "
          f"{len(equivalent_pairs)} equivalent-form positive controls.")

    formats = ("short", "medium", "long")

    # results[model][format][pair_id] = similarity
    results: dict[str, dict[str, dict[str, float]]] = {}
    # equivalent_results[model][pair_id] = similarity
    equivalent_results: dict[str, dict[str, float]] = {}

    for model_name, hf_id in MODELS.items():
        print(f"\nLoading {model_name} ({hf_id})...")
        model = SentenceTransformer(hf_id)
        results[model_name] = {f: {} for f in formats}
        equivalent_results[model_name] = {}

        # Distinct pairs across three formats
        for fmt in formats:
            method_name = {"short": "short", "medium": "medium", "long": "long_form"}[fmt]
            print(f"  Encoding distinct pairs (format={fmt})...")
            texts_a = [getattr(p, method_name)("a") for p in distinct_pairs]
            texts_b = [getattr(p, method_name)("b") for p in distinct_pairs]
            embs_a = encode_batch(model, texts_a)
            embs_b = encode_batch(model, texts_b)
            for p, ea, eb in zip(distinct_pairs, embs_a, embs_b):
                results[model_name][fmt][p.pair_id] = cosine_sim(ea, eb)

        # Equivalent-form pairs (positive control)
        print(f"  Encoding equivalent-form pairs...")
        eq_texts_a = [p.text_a() for p in equivalent_pairs]
        eq_texts_b = [p.text_b() for p in equivalent_pairs]
        eq_embs_a = encode_batch(model, eq_texts_a)
        eq_embs_b = encode_batch(model, eq_texts_b)
        for p, ea, eb in zip(equivalent_pairs, eq_embs_a, eq_embs_b):
            equivalent_results[model_name][p.pair_id] = cosine_sim(ea, eb)

        # Free model from memory before loading next
        del model

    # -----------------------------------------------------------------------
    # Output: full results.json
    # -----------------------------------------------------------------------
    output = {
        "models": list(MODELS.keys()),
        "formats": list(formats),
        "thresholds": list(THRESHOLDS),
        "n_distinct_pairs": len(distinct_pairs),
        "n_equivalent_pairs": len(equivalent_pairs),
        "distinct_pairs": [asdict(p) for p in distinct_pairs],
        "equivalent_pairs": [asdict(p) for p in equivalent_pairs],
        "similarities_distinct": results,
        "similarities_equivalent": equivalent_results,
    }
    (ROOT / "results.json").write_text(json.dumps(output, indent=2))
    print(f"\nWrote {ROOT / 'results.json'}")

    # -----------------------------------------------------------------------
    # Output: Table 1 — conflation rate by model x threshold (averaged across formats)
    # -----------------------------------------------------------------------
    n_distinct = len(distinct_pairs)
    rows = []
    for model_name in MODELS:
        # Average similarity across formats per pair, then threshold
        per_pair_avg = {
            p.pair_id: np.mean([results[model_name][f][p.pair_id] for f in formats])
            for p in distinct_pairs
        }
        row = {"model": model_name}
        for tau in THRESHOLDS:
            n_above = sum(1 for s in per_pair_avg.values() if s >= tau)
            row[f"tau_{tau}"] = round(n_above / n_distinct, 3)
        rows.append(row)
    rows.append({
        "model": "Typed-graph baseline",
        **{f"tau_{tau}": 0.0 for tau in THRESHOLDS},
    })
    with (ROOT / "conflation-rates.csv").open("w", newline="") as f:
        writer = csv.DictWriter(f, fieldnames=list(rows[0].keys()))
        writer.writeheader()
        writer.writerows(rows)
    print(f"Wrote {ROOT / 'conflation-rates.csv'}  (Table 1)")
    print()
    print("Table 1 (conflation rate at each threshold, averaged across formats):")
    for r in rows:
        print(f"  {r['model']:>22}  " + "  ".join(
            f"τ={tau}:{r[f'tau_{tau}']:.2f}" for tau in THRESHOLDS
        ))

    # -----------------------------------------------------------------------
    # Output: Table 2 — conflation rate at τ=0.95 by format x model
    # -----------------------------------------------------------------------
    fmt_rows = []
    for fmt in formats:
        row = {"format": fmt}
        for model_name in MODELS:
            sims = list(results[model_name][fmt].values())
            n_above = sum(1 for s in sims if s >= 0.95)
            row[model_name] = round(n_above / len(sims), 3)
        fmt_rows.append(row)
    with (ROOT / "format-rates.csv").open("w", newline="") as f:
        writer = csv.DictWriter(f, fieldnames=list(fmt_rows[0].keys()))
        writer.writeheader()
        writer.writerows(fmt_rows)
    print(f"\nWrote {ROOT / 'format-rates.csv'}  (Table 2 at τ=0.95)")
    print("\nTable 2 (conflation rate at τ=0.95 by format):")
    for r in fmt_rows:
        print(f"  {r['format']:>8}: " + "  ".join(
            f"{m}:{r[m]:.2f}" for m in MODELS
        ))

    # -----------------------------------------------------------------------
    # Output: Positive-control summary
    # -----------------------------------------------------------------------
    pc_summary = {}
    for model_name in MODELS:
        sims = list(equivalent_results[model_name].values())
        pc_summary[model_name] = {
            "n": len(sims),
            "mean": round(float(np.mean(sims)), 4),
            "sd": round(float(np.std(sims, ddof=1)), 4),
            "min": round(min(sims), 4),
            "max": round(max(sims), 4),
        }
    (ROOT / "positive-control.json").write_text(json.dumps(pc_summary, indent=2))
    print(f"\nPositive-control (equivalent-form pair similarities):")
    for m, stats in pc_summary.items():
        print(f"  {m}: mean={stats['mean']:.4f} sd={stats['sd']:.4f} "
              f"range=[{stats['min']:.4f}, {stats['max']:.4f}]")

    # -----------------------------------------------------------------------
    # Output: Figure 1 — per-pair, per-model heatmap
    # -----------------------------------------------------------------------
    if HAS_MPL:
        # Use medium-format similarities for the figure (most representative)
        labels = [f"{p.gene} {p.variant_a} vs {p.variant_b}" for p in distinct_pairs]
        matrix = np.array([
            [results[m]["medium"][p.pair_id] for m in MODELS]
            for p in distinct_pairs
        ])

        fig, ax = plt.subplots(figsize=(8, 0.45 * len(distinct_pairs) + 1.2))
        im = ax.imshow(matrix, aspect="auto", vmin=0.5, vmax=1.0, cmap="YlOrRd")
        ax.set_xticks(range(len(MODELS)))
        ax.set_xticklabels(list(MODELS.keys()), rotation=20, ha="right")
        ax.set_yticks(range(len(labels)))
        ax.set_yticklabels(labels, fontsize=8)
        ax.set_title("Cosine similarity: clinically distinct variant pairs\n"
                     "(medium format; higher = more conflation risk)",
                     fontsize=10)
        for i in range(matrix.shape[0]):
            for j in range(matrix.shape[1]):
                ax.text(j, i, f"{matrix[i,j]:.2f}",
                        ha="center", va="center", fontsize=7,
                        color="black" if matrix[i,j] < 0.85 else "white")
        cbar = fig.colorbar(im, ax=ax, fraction=0.04, pad=0.02)
        cbar.set_label("cosine similarity")
        ax.axhline(-0.5, color="black", linewidth=0.5)
        plt.tight_layout()
        plt.savefig(ROOT / "figure-1.png", dpi=200)
        print(f"\nWrote {ROOT / 'figure-1.png'}  (Figure 1)")

    print("\nDone. Drop the values from conflation-rates.csv and format-rates.csv "
          "into Section 4 of manuscript.md, paste positive-control.json values "
          "into Section 4.4, and use figure-1.png as Figure 1.")


if __name__ == "__main__":
    main()
```
