## Supplementary code and data: benchmark, experiment scripts, and result files for "Cosine Similarity Conflates Clinically Distinct Cancer Variants: A Case for Typed-Graph Retrieval in Precision Oncology Decision Support": experiment_asymmetric_long_format.py

```
#!/usr/bin/env python3
"""
experiment_asymmetric_long_format.py — Experiment 2.4: asymmetric long format.

The original long format (experiment.py) appended a single clinical-context
sentence that, because it described the contrast between the two variants,
was shared across both sides of a pair and trivially inflated cosine
similarity. This script re-runs the long-format measurement with a distinct,
variant-specific clinical sentence for each variant (no shared text). Each
sentence names the therapy or resistance pattern unique to that variant.

It also computes the lexical (Jaccard) overlap between each pair's two
sentences, to confirm that the experiment isolates the clinical-context
effect from raw token overlap.

Outputs:
  asymmetric-long-format-rates.csv   conflation rate per threshold x model
                                     (same column schema as format-rates.csv)
  asymmetric-long-per-pair.csv       pair_id, model, cosine_similarity
  asymmetric-long-jaccard.csv        pair_id, gene, jaccard, shared/union tokens

Runtime: ~1 minute (loads up to three embedding models on first run).
"""
from __future__ import annotations

import re

from experiment_common import (ROOT, MODELS, THRESHOLDS, VARIANTS, embed, cosine,
                                load_pairs, variant_key, write_csv)


def tokens(text: str) -> set[str]:
    return {t for t in re.split(r"[^a-z0-9]+", text.lower()) if t}


def main() -> None:
    distinct, _ = load_pairs()

    # Per-pair cosine in the asymmetric (variant-specific) long format.
    per_pair_rows = []
    sims: dict[str, list[float]] = {m: [] for m in MODELS}
    for model in MODELS:
        texts_a = [VARIANTS[variant_key(p.gene, p.variant_a)]["asym"] for p in distinct]
        texts_b = [VARIANTS[variant_key(p.gene, p.variant_b)]["asym"] for p in distinct]
        ea, eb = embed(model, texts_a), embed(model, texts_b)
        for p, va, vb in zip(distinct, ea, eb):
            s = cosine(va, vb)
            sims[model].append(s)
            per_pair_rows.append(dict(pair_id=p.pair_id, model=model,
                                      cosine_similarity=round(s, 4)))
    write_csv(ROOT / "asymmetric-long-per-pair.csv",
              ["pair_id", "model", "cosine_similarity"], per_pair_rows)

    # Conflation rates per threshold, same column schema as format-rates.csv.
    rate_rows = []
    for tau in THRESHOLDS:
        row = {"format": f"asymmetric-long @ tau={tau}"}
        for model in MODELS:
            vals = sims[model]
            row[model] = round(sum(1 for s in vals if s >= tau) / len(vals), 4)
        rate_rows.append(row)
    write_csv(ROOT / "asymmetric-long-format-rates.csv",
              ["format"] + list(MODELS), rate_rows)

    # Lexical overlap between the two sentences of each pair.
    jac_rows = []
    for p in distinct:
        ta = tokens(VARIANTS[variant_key(p.gene, p.variant_a)]["asym"])
        tb = tokens(VARIANTS[variant_key(p.gene, p.variant_b)]["asym"])
        inter, union = ta & tb, ta | tb
        jac_rows.append(dict(pair_id=p.pair_id, gene=p.gene,
                             jaccard=round(len(inter) / len(union), 4),
                             shared_tokens=len(inter), union_tokens=len(union)))
    write_csv(ROOT / "asymmetric-long-jaccard.csv",
              ["pair_id", "gene", "jaccard", "shared_tokens", "union_tokens"], jac_rows)

    mean_jac = sum(r["jaccard"] for r in jac_rows) / len(jac_rows)
    print(f"  mean pairwise Jaccard overlap: {mean_jac:.3f}")
    for model in MODELS:
        vals = sims[model]
        print(f"  {model:>20}: asymmetric-long cosine min={min(vals):.3f} "
              f"max={max(vals):.3f} mean={sum(vals)/len(vals):.3f}")


if __name__ == "__main__":
    main()
```
