## Supplementary code and data: benchmark, experiment scripts, and result files for "Cosine Similarity Conflates Clinically Distinct Cancer Variants: A Case for Typed-Graph Retrieval in Precision Oncology Decision Support": experiment_ci.py

```
#!/usr/bin/env python3
"""
experiment_ci.py — Experiment 2.1: exact binomial 95% confidence intervals.

With nine variant pairs, every reported conflation proportion needs an
interval. This script:

  * reads the per-pair similarities in results.json (and reproduces
    conflation-rates.csv) and attaches an exact (Clopper-Pearson) binomial
    95% CI to every conflation proportion, per model x format x threshold;
  * bootstraps a 95% CI (10,000 resamples) for the equivalent-form
    positive-control mean similarity, per encoder.

Outputs:
  conflation-rates-with-ci.csv   model, threshold, format, n, k, proportion,
                                 ci_lower, ci_upper
  positive-control-with-ci.json  mean, sd, ci_lower, ci_upper per encoder

No embedding models are loaded; this script reads existing outputs only.
Runtime: < 5 seconds.
"""
from __future__ import annotations

import json

import numpy as np
from scipy.stats import binomtest

from experiment_common import ROOT, THRESHOLDS, MODELS, load_results, write_csv

RNG = np.random.default_rng(20260505)
N_BOOT = 10_000


def binom_ci(k: int, n: int) -> tuple[float, float]:
    ci = binomtest(k, n).proportion_ci(method="exact", confidence_level=0.95)
    return round(ci.low, 4), round(ci.high, 4)


def main() -> None:
    res = load_results()
    formats = res["formats"]
    pairs = [p["pair_id"] for p in res["distinct_pairs"]]
    n = len(pairs)
    sims = res["similarities_distinct"]

    rows = []
    # Per-format conflation rates with CIs.
    for model in MODELS:
        for fmt in formats:
            for tau in THRESHOLDS:
                k = sum(1 for pid in pairs if sims[model][fmt][pid] >= tau)
                lo, hi = binom_ci(k, n)
                rows.append(dict(model=model, threshold=tau, format=fmt, n=n, k=k,
                                 proportion=round(k / n, 4), ci_lower=lo, ci_upper=hi))
    # Format-averaged conflation rates with CIs (reproduces conflation-rates.csv).
    for model in MODELS:
        per_pair_avg = {pid: float(np.mean([sims[model][f][pid] for f in formats]))
                        for pid in pairs}
        for tau in THRESHOLDS:
            k = sum(1 for pid in pairs if per_pair_avg[pid] >= tau)
            lo, hi = binom_ci(k, n)
            rows.append(dict(model=model, threshold=tau, format="averaged", n=n, k=k,
                             proportion=round(k / n, 4), ci_lower=lo, ci_upper=hi))
    # Typed-graph baseline: conflation is zero by construction. The CI is the
    # exact interval for an observed 0/9 and is reported for completeness.
    for tau in THRESHOLDS:
        lo, hi = binom_ci(0, n)
        rows.append(dict(model="Typed-graph baseline", threshold=tau, format="averaged",
                         n=n, k=0, proportion=0.0, ci_lower=lo, ci_upper=hi))

    write_csv(ROOT / "conflation-rates-with-ci.csv",
              ["model", "threshold", "format", "n", "k", "proportion",
               "ci_lower", "ci_upper"], rows)

    # Bootstrap CIs for the equivalent-form positive-control means.
    pc = {}
    for model in MODELS:
        vals = np.array(list(res["similarities_equivalent"][model].values()), dtype=float)
        boot = np.array([RNG.choice(vals, size=len(vals), replace=True).mean()
                         for _ in range(N_BOOT)])
        lo, hi = np.percentile(boot, [2.5, 97.5])
        pc[model] = dict(n=len(vals), mean=round(float(vals.mean()), 4),
                         sd=round(float(vals.std(ddof=1)), 4),
                         ci_lower=round(float(lo), 4), ci_upper=round(float(hi), 4),
                         n_bootstrap=N_BOOT)
    (ROOT / "positive-control-with-ci.json").write_text(json.dumps(pc, indent=2))
    print("Wrote positive-control-with-ci.json")
    for m, s in pc.items():
        print(f"  {m}: mean={s['mean']} 95% CI [{s['ci_lower']}, {s['ci_upper']}]")


if __name__ == "__main__":
    main()
```
