## Supplementary code and data: benchmark, experiment scripts, and result files for "Cosine Similarity Conflates Clinically Distinct Cancer Variants: A Case for Typed-Graph Retrieval in Precision Oncology Decision Support": experiment_common.py

```
#!/usr/bin/env python3
"""
experiment_common.py — shared helpers for the v2 revision experiments.

The v2 revision adds eight new experiment scripts (experiment_ci.py,
experiment_negative_controls.py, experiment_separation.py,
experiment_asymmetric_long_format.py, experiment_corpus_retrieval.py,
experiment_id_guardrail.py, experiment_typed_graph.py,
experiment_loo_sensitivity.py). This module holds the data, the embedding
cache, and the small utilities they share so that:

  * each script stays self-contained and individually runnable;
  * embeddings are computed once and reused across scripts (disk cache in
    `.emb-cache/`), keeping the full v2 re-run well under 30 minutes;
  * variant text rendering is identical to the original `experiment.py`.

No external API dependencies. Open-source embedding models only.
Delete `.emb-cache/` to force a clean recomputation.
"""
from __future__ import annotations

import csv
import json
from pathlib import Path
from typing import Sequence

import numpy as np

# Reuse the original variant rendering / loading logic verbatim so the v2
# experiments operate on exactly the same text the v1 benchmark used.
from experiment import VariantPair, EquivalentPair, load_pairs  # noqa: F401

ROOT = Path(__file__).resolve().parent
CACHE_DIR = ROOT / ".emb-cache"
THRESHOLDS = (0.85, 0.90, 0.95, 0.99)

MODELS = {
    "PubMedBERT": "microsoft/BiomedNLP-BiomedBERT-base-uncased-abstract-fulltext",
    "MedCPT": "ncbi/MedCPT-Article-Encoder",
    "BGE-large-en-v1.5": "BAAI/bge-large-en-v1.5",
}
BIOMEDICAL_MODELS = ("PubMedBERT", "MedCPT")

# ---------------------------------------------------------------------------
# Embedding cache. Embeddings are deterministic given (model, text), so we
# cache them keyed by text. The first v2 script that needs a model loads it;
# later scripts reuse the disk cache and never re-load a model.
# ---------------------------------------------------------------------------

_loaded: dict[str, object] = {}
_disk: dict[str, dict[str, list]] = {}


def _cache_path(model_name: str) -> Path:
    return CACHE_DIR / f"{model_name.replace('/', '_')}.json"


def _disk_cache(model_name: str) -> dict[str, list]:
    if model_name not in _disk:
        p = _cache_path(model_name)
        _disk[model_name] = json.loads(p.read_text()) if p.exists() else {}
    return _disk[model_name]


def embed(model_name: str, texts: Sequence[str]) -> np.ndarray:
    """Return an (len(texts), dim) array of L2-normalized embeddings."""
    texts = list(texts)
    cache = _disk_cache(model_name)
    missing = sorted({t for t in texts if t not in cache})
    if missing:
        from sentence_transformers import SentenceTransformer

        if model_name not in _loaded:
            print(f"  [loading embedding model: {model_name}]")
            _loaded[model_name] = SentenceTransformer(MODELS[model_name])
        model = _loaded[model_name]
        vecs = model.encode(missing, normalize_embeddings=True, show_progress_bar=False)
        for t, v in zip(missing, vecs):
            cache[t] = [float(x) for x in v]
        CACHE_DIR.mkdir(exist_ok=True)
        _cache_path(model_name).write_text(json.dumps(cache))
    return np.array([cache[t] for t in texts], dtype=float)


def cosine(a: np.ndarray, b: np.ndarray) -> float:
    a = np.asarray(a, dtype=float)
    b = np.asarray(b, dtype=float)
    return float(np.dot(a, b) / ((np.linalg.norm(a) + 1e-12) * (np.linalg.norm(b) + 1e-12)))


def load_results() -> dict:
    """Return the parsed results.json produced by the v1 experiment.py."""
    return json.loads((ROOT / "results.json").read_text())


def write_csv(path: Path, fieldnames: Sequence[str], rows: Sequence[dict]) -> None:
    with path.open("w", newline="") as f:
        w = csv.DictWriter(f, fieldnames=list(fieldnames))
        w.writeheader()
        w.writerows(rows)
    print(f"Wrote {path.name}  ({len(rows)} rows)")


# ---------------------------------------------------------------------------
# Canonical variant table — single source of truth for the v2 experiments
# that need a knowledge base (corpus retrieval, ID guardrail, typed graph).
# Sixteen distinct variants appear across the nine clinically distinct pairs
# (EGFR L858R and KRAS G12C each appear in two pairs). `hgvs_protein` is the
# canonical node identifier; `drugs` / `tier` populate the typed graph.
# Therapy associations and AMP/ASCO/CAP tiers follow the variant-pair set and
# the FDA-approval landscape cited in the manuscript.
# ---------------------------------------------------------------------------

VARIANTS: dict[str, dict] = {
    "BRAF V600E": dict(
        gene="BRAF", variant_raw="V600E", hgvs_protein="V600E", tumor="melanoma",
        short="BRAF V600E", drugs=["dabrafenib + trametinib"], tier="I-A",
        asym="BRAF V600E in melanoma; targetable by combined dabrafenib plus trametinib, "
             "with the highest response rates among BRAF-mutant melanoma in the COMBI-d and COMBI-v trials.",
    ),
    "BRAF V600K": dict(
        gene="BRAF", variant_raw="V600K", hgvs_protein="V600K", tumor="melanoma",
        short="BRAF V600K", drugs=["dabrafenib + trametinib"], tier="I-A",
        asym="BRAF V600K in melanoma; responds to combined BRAF and MEK inhibition but with "
             "lower response rates and shorter progression-free survival than V600E.",
    ),
    "EGFR L858R": dict(
        gene="EGFR", variant_raw="L858R", hgvs_protein="L858R", tumor="non-small cell lung cancer",
        short="EGFR L858R", drugs=["osimertinib"], tier="I-A",
        asym="EGFR L858R in non-small cell lung cancer; a sensitizing kinase-domain mutation for "
             "which osimertinib is first-line standard of care per the FLAURA trial.",
    ),
    "EGFR T790M": dict(
        gene="EGFR", variant_raw="T790M", hgvs_protein="T790M", tumor="non-small cell lung cancer",
        short="EGFR T790M", drugs=["osimertinib"], tier="I-A",
        asym="EGFR T790M in non-small cell lung cancer; the gatekeeper resistance mutation that "
             "emerges on early-generation tyrosine kinase inhibitors and indicates second-line osimertinib per AURA3.",
    ),
    "EGFR exon-19-deletion": dict(
        gene="EGFR", variant_raw="exon-19-deletion", hgvs_protein="exon19del",
        tumor="non-small cell lung cancer", short="EGFR exon 19 deletion",
        drugs=["osimertinib"], tier="I-A",
        asym="EGFR exon 19 deletion in non-small cell lung cancer; a sensitizing alteration with "
             "higher response rates and longer progression-free survival on osimertinib than other EGFR variants.",
    ),
    "KRAS G12C": dict(
        gene="KRAS", variant_raw="G12C", hgvs_protein="G12C", tumor="non-small cell lung cancer",
        short="KRAS G12C", drugs=["sotorasib", "adagrasib"], tier="I-A",
        asym="KRAS G12C in non-small cell lung cancer; targetable by the covalent inhibitors "
             "sotorasib and adagrasib, which depend on the mutant cysteine residue at codon 12.",
    ),
    "KRAS G12D": dict(
        gene="KRAS", variant_raw="G12D", hgvs_protein="G12D", tumor="non-small cell lung cancer",
        short="KRAS G12D", drugs=[], tier="III",
        asym="KRAS G12D in non-small cell lung cancer; has no FDA-approved targeted therapy and "
             "requires a distinct noncovalent inhibitor chemotype such as MRTX1133.",
    ),
    "KRAS G12V": dict(
        gene="KRAS", variant_raw="G12V", hgvs_protein="G12V", tumor="non-small cell lung cancer",
        short="KRAS G12V", drugs=[], tier="III",
        asym="KRAS G12V in non-small cell lung cancer; lacks an FDA-approved targeted therapy and "
             "is currently addressed only by investigational pan-RAS inhibitors.",
    ),
    "ERBB2 amplification": dict(
        gene="ERBB2", variant_raw="amplification", hgvs_protein="amplification",
        tumor="non-small cell lung cancer", short="ERBB2 amplification",
        drugs=["trastuzumab"], tier="I-A",
        asym="ERBB2 amplification; a copy-number alteration targeted by trastuzumab-based "
             "HER2-directed antibody therapy.",
    ),
    "ERBB2 G776delinsVC": dict(
        gene="ERBB2", variant_raw="G776delinsVC", hgvs_protein="G776delinsVC",
        tumor="non-small cell lung cancer", short="ERBB2 G776delinsVC",
        drugs=["trastuzumab deruxtecan"], tier="I-B",
        asym="ERBB2 G776delinsVC, an exon 20 activating insertion in non-small cell lung cancer; "
             "targeted by trastuzumab deruxtecan per the DESTINY-Lung02 trial.",
    ),
    "PIK3CA E545K": dict(
        gene="PIK3CA", variant_raw="E545K", hgvs_protein="E545K", tumor="breast cancer",
        short="PIK3CA E545K", drugs=["alpelisib"], tier="I-A",
        asym="PIK3CA E545K in breast cancer; an activating helical-domain mutation on the SOLAR-1 "
             "alpelisib and INAVO120 inavolisib eligibility lists.",
    ),
    "PIK3CA M1043I": dict(
        gene="PIK3CA", variant_raw="M1043I", hgvs_protein="M1043I", tumor="breast cancer",
        short="PIK3CA M1043I", drugs=["alpelisib"], tier="II-D",
        asym="PIK3CA M1043I in breast cancer; an activating mutation supported by lower-tier "
             "evidence and outside the original SOLAR-1 alpelisib eligibility set.",
    ),
    "PIK3CA H1047R": dict(
        gene="PIK3CA", variant_raw="H1047R", hgvs_protein="H1047R", tumor="breast cancer",
        short="PIK3CA H1047R", drugs=["alpelisib"], tier="I-A",
        asym="PIK3CA H1047R in breast cancer; a kinase-domain hotspot with allele-specific "
             "SOLAR-1 alpelisib hazard ratios.",
    ),
    "PIK3CA E542K": dict(
        gene="PIK3CA", variant_raw="E542K", hgvs_protein="E542K", tumor="breast cancer",
        short="PIK3CA E542K", drugs=["alpelisib"], tier="I-A",
        asym="PIK3CA E542K in breast cancer; a helical-domain hotspot with SOLAR-1 hazard ratios "
             "that differ from other PIK3CA hotspots.",
    ),
    "NTRK1 fusion": dict(
        gene="NTRK1", variant_raw="fusion", hgvs_protein="fusion", tumor="solid tumor",
        short="NTRK1 fusion", drugs=["larotrectinib", "entrectinib"], tier="I-A",
        asym="NTRK1 gene fusion; a tumor-agnostic alteration with FDA approval for larotrectinib "
             "and entrectinib.",
    ),
    "NTRK1 point-mutation": dict(
        gene="NTRK1", variant_raw="point-mutation", hgvs_protein="G595R",
        tumor="non-small cell lung cancer", short="NTRK1 point mutation",
        drugs=[], tier="III",
        asym="NTRK1 G595R, a solvent-front kinase-domain point mutation; confers resistance to "
             "first-generation TRK inhibitors and is not independently FDA-targetable.",
    ),
}


def variant_key(gene: str, variant_raw: str) -> str:
    """Map a (gene, raw variant token) from variant-pairs.csv to a VARIANTS key."""
    return f"{gene} {variant_raw}"


# ---------------------------------------------------------------------------
# Cross-gene negative-control decoys (Experiment 2.2, Set A). Nine canonical
# cancer driver variants, each from a different gene and tumor type than the
# main-set variant it is paired with. Template-matched ("gene variant in
# tumor"), but biologically unrelated.
# ---------------------------------------------------------------------------

DECOYS_SET_A: list[dict] = [
    dict(gene="TP53", variant="R175H", hgvs_protein="R175H", tumor="colorectal cancer",
         context="TP53 R175H in colorectal cancer; a structural hotspot mutation in the p53 tumor suppressor."),
    dict(gene="APC", variant="R1450X", hgvs_protein="R1450X", tumor="colorectal cancer",
         context="APC R1450X in colorectal cancer; a truncating mutation in the APC tumor suppressor."),
    dict(gene="RB1", variant="R661W", hgvs_protein="R661W", tumor="retinoblastoma",
         context="RB1 R661W in retinoblastoma; a mutation inactivating the RB1 tumor suppressor."),
    dict(gene="VHL", variant="L188V", hgvs_protein="L188V", tumor="renal cell carcinoma",
         context="VHL L188V in renal cell carcinoma; a mutation in the von Hippel-Lindau tumor suppressor."),
    dict(gene="PTEN", variant="R130G", hgvs_protein="R130G", tumor="endometrial carcinoma",
         context="PTEN R130G in endometrial carcinoma; a phosphatase-domain mutation inactivating PTEN."),
    dict(gene="MYC", variant="amplification", hgvs_protein="amplification", tumor="neuroblastoma",
         context="MYC amplification in neuroblastoma; a copy-number gain of the MYC oncogene."),
    dict(gene="MLH1", variant="V506A", hgvs_protein="V506A", tumor="colorectal cancer",
         context="MLH1 V506A in colorectal cancer; a mismatch-repair gene variant relevant to Lynch syndrome."),
    dict(gene="CTNNB1", variant="S37F", hgvs_protein="S37F", tumor="hepatocellular carcinoma",
         context="CTNNB1 S37F in hepatocellular carcinoma; a beta-catenin exon 3 activating mutation."),
    dict(gene="SMAD4", variant="R361H", hgvs_protein="R361H", tumor="pancreatic cancer",
         context="SMAD4 R361H in pancreatic cancer; a mutation in the SMAD4 tumor suppressor."),
]

# Same-gene, non-canonical tumor type (Experiment 2.2, Set B). The tumor type
# paired against each main-set variant_a to probe tumor-type sensitivity.
SAME_GENE_OTHER_TUMOR: dict[str, str] = {
    "BRAF V600E": "colorectal cancer",
    "EGFR L858R": "glioblastoma",
    "EGFR exon-19-deletion": "glioblastoma",
    "KRAS G12C": "pancreatic cancer",
    "ERBB2 amplification": "gastric cancer",
    "PIK3CA E545K": "endometrial carcinoma",
    "PIK3CA H1047R": "head and neck carcinoma",
    "NTRK1 fusion": "colorectal cancer",
}

# Thirty cross-gene hard-negative decoys for the retrieval corpus
# (Experiments 2.5 / 2.6). Canonical variants from thirty highly cited cancer
# genes, none of which is one of the six main-set genes, all in the
# "gene variant in tumor type" template.
CORPUS_DECOYS: list[dict] = [
    dict(gene="TP53", variant="R175H", hgvs_protein="R175H", tumor="colorectal cancer"),
    dict(gene="APC", variant="R1450X", hgvs_protein="R1450X", tumor="colorectal cancer"),
    dict(gene="PTEN", variant="R130G", hgvs_protein="R130G", tumor="endometrial carcinoma"),
    dict(gene="RB1", variant="R661W", hgvs_protein="R661W", tumor="retinoblastoma"),
    dict(gene="VHL", variant="L188V", hgvs_protein="L188V", tumor="renal cell carcinoma"),
    dict(gene="CTNNB1", variant="S37F", hgvs_protein="S37F", tumor="hepatocellular carcinoma"),
    dict(gene="SMAD4", variant="R361H", hgvs_protein="R361H", tumor="pancreatic cancer"),
    dict(gene="MLH1", variant="V506A", hgvs_protein="V506A", tumor="colorectal cancer"),
    dict(gene="IDH1", variant="R132H", hgvs_protein="R132H", tumor="glioma"),
    dict(gene="IDH2", variant="R140Q", hgvs_protein="R140Q", tumor="acute myeloid leukemia"),
    dict(gene="FLT3", variant="D835Y", hgvs_protein="D835Y", tumor="acute myeloid leukemia"),
    dict(gene="NPM1", variant="W288fs", hgvs_protein="W288fs", tumor="acute myeloid leukemia"),
    dict(gene="JAK2", variant="V617F", hgvs_protein="V617F", tumor="myeloproliferative neoplasm"),
    dict(gene="KIT", variant="D816V", hgvs_protein="D816V", tumor="systemic mastocytosis"),
    dict(gene="PDGFRA", variant="D842V", hgvs_protein="D842V", tumor="gastrointestinal stromal tumor"),
    dict(gene="MET", variant="exon-14-skipping", hgvs_protein="exon14skip", tumor="non-small cell lung cancer"),
    dict(gene="ALK", variant="F1174L", hgvs_protein="F1174L", tumor="neuroblastoma"),
    dict(gene="ROS1", variant="G2032R", hgvs_protein="G2032R", tumor="non-small cell lung cancer"),
    dict(gene="RET", variant="M918T", hgvs_protein="M918T", tumor="medullary thyroid carcinoma"),
    dict(gene="FGFR3", variant="S249C", hgvs_protein="S249C", tumor="bladder cancer"),
    dict(gene="AKT1", variant="E17K", hgvs_protein="E17K", tumor="breast cancer"),
    dict(gene="ESR1", variant="Y537S", hgvs_protein="Y537S", tumor="breast cancer"),
    dict(gene="NRAS", variant="Q61K", hgvs_protein="Q61K", tumor="melanoma"),
    dict(gene="HRAS", variant="G13R", hgvs_protein="G13R", tumor="head and neck carcinoma"),
    dict(gene="CDKN2A", variant="R80X", hgvs_protein="R80X", tumor="pancreatic cancer"),
    dict(gene="ATM", variant="R337H", hgvs_protein="R337H", tumor="breast cancer"),
    dict(gene="BRCA1", variant="C61G", hgvs_protein="C61G", tumor="ovarian cancer"),
    dict(gene="BRCA2", variant="N3124I", hgvs_protein="N3124I", tumor="ovarian cancer"),
    dict(gene="POLE", variant="P286R", hgvs_protein="P286R", tumor="endometrial carcinoma"),
    dict(gene="STK11", variant="F354L", hgvs_protein="F354L", tumor="non-small cell lung cancer"),
]


# ---------------------------------------------------------------------------
# Ingestion-time variant-string normalization (Experiments 2.6 and 2.7).
# Resolves a free-text variant string to a canonical (gene, hgvs_protein)
# tuple, or None when the string is too underspecified to resolve. The rules
# are general regex patterns for the standard variant classes; they are NOT
# hand-tuned to the benchmark, so non-point-mutation classes are characterized
# honestly.
# ---------------------------------------------------------------------------

import re

_GENE_RE = re.compile(r"^([A-Z][A-Z0-9]{1,9})\b")
_POINT_RE = re.compile(r"\b([A-Z]\d{1,4}[A-Z])\b")
_DELINS_RE = re.compile(r"\b([A-Z]\d{1,4}delins[A-Z]+)\b")
_FS_RE = re.compile(r"\b([A-Z]\d{1,4}fs)\b")
_EXON_RE = re.compile(r"\bexon[\s-]*(\d{1,2})[\s-]*(deletion|insertion|skipping|del|ins|skip)\b", re.I)
_AMP_RE = re.compile(r"\bamplif", re.I)
_FUSION_RE = re.compile(r"\bfusion\b", re.I)


def normalize_variant_string(text: str) -> tuple[str, str] | None:
    """Resolve a variant string to a canonical (gene, hgvs_protein) tuple.

    Returns None when the string carries no resolvable variant token (e.g. a
    bare "point mutation" with no residue specified). Only the leading
    gene-anchored segment is parsed, so trailing clinical context does not
    contaminate the resolution.
    """
    text = text.strip()
    gm = _GENE_RE.match(text)
    if not gm:
        return None
    gene = gm.group(1)
    # Restrict the variant search to the segment before " in <tumor>" / ";".
    head = re.split(r"\s+in\s+|;", text, maxsplit=1)[0]
    remainder = head[gm.end():]

    m = _DELINS_RE.search(remainder)
    if m:
        return (gene, m.group(1))
    m = _FS_RE.search(remainder)
    if m:
        return (gene, m.group(1))
    m = _EXON_RE.search(remainder)
    if m:
        kind = "skip" if m.group(2).lower().startswith("skip") else (
            "del" if m.group(2).lower().startswith("del") else "ins")
        return (gene, f"exon{m.group(1)}{kind}")
    m = _POINT_RE.search(remainder)
    if m:
        return (gene, m.group(1))
    if _AMP_RE.search(remainder):
        return (gene, "amplification")
    if _FUSION_RE.search(remainder):
        return (gene, "fusion")
    return None


# ---------------------------------------------------------------------------
# Retrieval corpus and ranked-retrieval metrics (Experiments 2.5 and 2.6).
# The corpus is shared by both scripts so the ID-guardrail experiment runs
# over exactly the same documents as the plain cosine-retrieval experiment.
# ---------------------------------------------------------------------------

import math


def build_corpus() -> list[dict]:
    """Build the retrieval corpus: 16 canonical variant docs + 6
    equivalent-notation docs + 30 cross-gene hard-negative decoys = 52 docs.

    Each doc carries a `variant_id` = (gene, hgvs_protein). Equivalent-notation
    docs share the canonical variant_id of the variant they re-notate.
    """
    corpus: list[dict] = []
    for key, v in VARIANTS.items():
        corpus.append(dict(doc_id=f"canon:{key}", kind="canonical",
                           variant_id=(v["gene"], v["hgvs_protein"]),
                           text=f"{v['short']} in {v['tumor']}"))
    # Equivalent-notation docs from equivalent-pairs.csv.
    _, equivalent = load_pairs()
    for e in equivalent:
        norm = normalize_variant_string(f"{e.gene} {e.variant_a}")
        text = e.variant_b if e.variant_b.startswith(f"{e.gene} ") else f"{e.gene} {e.variant_b}"
        corpus.append(dict(doc_id=f"equiv:{e.pair_id}", kind="equivalent",
                           variant_id=norm, text=text))
    for d in CORPUS_DECOYS:
        corpus.append(dict(doc_id=f"decoy:{d['gene']}:{d['hgvs_protein']}", kind="decoy",
                           variant_id=(d["gene"], d["hgvs_protein"]),
                           text=f"{d['gene']} {d['variant']} in {d['tumor']}"))
    return corpus


def build_queries() -> list[dict]:
    """Build the 16 retrieval queries (one per distinct main-set variant),
    each carrying its canonical variant_id and the set of partner variant_ids
    it is paired with across the nine clinically distinct pairs.
    """
    distinct, _ = load_pairs()
    partners: dict[str, set] = {k: set() for k in VARIANTS}
    for p in distinct:
        ka, kb = variant_key(p.gene, p.variant_a), variant_key(p.gene, p.variant_b)
        partners[ka].add((VARIANTS[kb]["gene"], VARIANTS[kb]["hgvs_protein"]))
        partners[kb].add((VARIANTS[ka]["gene"], VARIANTS[ka]["hgvs_protein"]))
    queries = []
    for key, v in VARIANTS.items():
        queries.append(dict(key=key, variant_id=(v["gene"], v["hgvs_protein"]),
                            partner_ids=partners[key],
                            texts={"short": v["short"],
                                   "medium": f"{v['short']} in {v['tumor']}",
                                   "long": v["asym"]}))
    return queries


def _dcg(rels: list[int]) -> float:
    return sum(r / math.log2(i + 2) for i, r in enumerate(rels))


def retrieval_metrics(ranked_ids: list, correct_id, partner_ids: set, k: int) -> dict:
    """Compute wrong_variant@k, recall@k, MRR, and nDCG@k for one query given
    a ranked list of doc variant_ids (best first)."""
    topk = ranked_ids[:k]
    recall = any(vid == correct_id for vid in topk)
    wrong = any(vid in partner_ids for vid in topk)
    mrr = 0.0
    for rank, vid in enumerate(ranked_ids, start=1):
        if vid == correct_id:
            mrr = 1.0 / rank
            break
    rels = [1 if vid == correct_id else 0 for vid in topk]
    n_correct_total = sum(1 for vid in ranked_ids if vid == correct_id)
    idcg = _dcg([1] * min(k, n_correct_total))
    ndcg = (_dcg(rels) / idcg) if idcg > 0 else 0.0
    return dict(recall=recall, wrong=wrong, mrr=mrr, ndcg=ndcg)
```
