## Supplementary code and data: benchmark, experiment scripts, and result files for "Cosine Similarity Conflates Clinically Distinct Cancer Variants: A Case for Typed-Graph Retrieval in Precision Oncology Decision Support": experiment_corpus_retrieval.py

```
#!/usr/bin/env python3
"""
experiment_corpus_retrieval.py — Experiment 2.5: corpus-level ranked retrieval.

Pairwise cosine similarity is a proxy; production RAG performs ranked
retrieval over a corpus. This script builds a 52-document corpus (16 canonical
variant docs + 6 equivalent-notation docs + 30 cross-gene hard-negative
decoys) and, for each of the 16 distinct main-set variants as a query,
retrieves the top-k by cosine similarity.

It reports, for k in {1, 3, 5}, averaged over the 16 queries:
  wrong_variant_at_k  fraction of queries where a paired (wrong) variant
                      appears in the top-k
  recall_at_k         fraction where the correct variant is retrieved
  mrr                 mean reciprocal rank of the correct variant
  ndcg_at_k           normalized discounted cumulative gain

Output:
  corpus-retrieval-metrics.csv   model, format, k, wrong_variant_at_k,
                                 recall_at_k, mrr, ndcg_at_k

Runtime: ~1-2 minutes (loads up to three embedding models on first run).
"""
from __future__ import annotations

import numpy as np

from experiment_common import (ROOT, MODELS, embed, build_corpus, build_queries,
                                retrieval_metrics, write_csv)

FORMATS = ("short", "medium", "long")
KS = (1, 3, 5)


def main() -> None:
    corpus = build_corpus()
    queries = build_queries()
    doc_ids = [d["variant_id"] for d in corpus]
    print(f"Corpus: {len(corpus)} documents; {len(queries)} queries.")

    rows = []
    for model in MODELS:
        doc_emb = embed(model, [d["text"] for d in corpus])
        for fmt in FORMATS:
            q_emb = embed(model, [q["texts"][fmt] for q in queries])
            # Per-query ranked variant_id lists.
            ranked_per_query = []
            for qi in range(len(queries)):
                scores = doc_emb @ q_emb[qi]
                order = np.argsort(-scores)
                ranked_per_query.append([doc_ids[i] for i in order])
            for k in KS:
                agg = {"wrong": [], "recall": [], "mrr": [], "ndcg": []}
                for q, ranked in zip(queries, ranked_per_query):
                    m = retrieval_metrics(ranked, q["variant_id"],
                                          q["partner_ids"], k)
                    agg["wrong"].append(m["wrong"])
                    agg["recall"].append(m["recall"])
                    agg["mrr"].append(m["mrr"])
                    agg["ndcg"].append(m["ndcg"])
                rows.append(dict(
                    model=model, format=fmt, k=k,
                    wrong_variant_at_k=round(float(np.mean(agg["wrong"])), 4),
                    recall_at_k=round(float(np.mean(agg["recall"])), 4),
                    mrr=round(float(np.mean(agg["mrr"])), 4),
                    ndcg_at_k=round(float(np.mean(agg["ndcg"])), 4),
                ))

    write_csv(ROOT / "corpus-retrieval-metrics.csv",
              ["model", "format", "k", "wrong_variant_at_k", "recall_at_k",
               "mrr", "ndcg_at_k"], rows)

    for r in rows:
        if r["format"] == "medium" and r["k"] == 5:
            print(f"  {r['model']:>20} (medium, k=5): "
                  f"wrong_variant@5={r['wrong_variant_at_k']}  "
                  f"recall@5={r['recall_at_k']}")


if __name__ == "__main__":
    main()
```
