## Supplementary code and data: benchmark, experiment scripts, and result files for "Cosine Similarity Conflates Clinically Distinct Cancer Variants: A Case for Typed-Graph Retrieval in Precision Oncology Decision Support": experiment_id_guardrail.py

```
#!/usr/bin/env python3
"""
experiment_id_guardrail.py — Experiment 2.6: exact-ID guardrail baseline.

A standard production hardening for vector retrieval is to follow cosine
retrieval with an exact variant-ID post-filter. This script implements that
two-stage pipeline over the same 52-document corpus and 16 queries as
Experiment 2.5:

  Stage 1: cosine retrieval (identical to experiment_corpus_retrieval.py)
  Stage 2: keep only top-k documents whose (gene, hgvs_protein) exactly
           matches the query's canonical ID, extracted from the query text
           by normalize_variant_string()

It reports the same metrics (wrong_variant_at_k, recall_at_k, mrr,
ndcg_at_k) so the guardrail can be compared directly against plain cosine
retrieval and against the typed-graph pipeline (Experiment 2.7).

Output:
  id-guardrail-metrics.csv   model, format, k, wrong_variant_at_k,
                             recall_at_k, mrr, ndcg_at_k

Runtime: < 1 minute; reuses cached embeddings from Experiment 2.5 when
available, so no embedding model is re-loaded if 2.5 ran first.
"""
from __future__ import annotations

import numpy as np

from experiment_common import (ROOT, MODELS, embed, build_corpus, build_queries,
                                retrieval_metrics, normalize_variant_string, write_csv)

FORMATS = ("short", "medium", "long")
KS = (1, 3, 5)


def main() -> None:
    corpus = build_corpus()
    queries = build_queries()
    doc_ids = [d["variant_id"] for d in corpus]
    print(f"Corpus: {len(corpus)} documents; {len(queries)} queries "
          f"(stage 1 cosine + stage 2 exact-ID filter).")

    rows = []
    for model in MODELS:
        doc_emb = embed(model, [d["text"] for d in corpus])
        for fmt in FORMATS:
            q_texts = [q["texts"][fmt] for q in queries]
            q_emb = embed(model, q_texts)
            # Stage 1 ranking, then stage 2 exact-ID filter.
            ranked_per_query = []
            for qi, q in enumerate(queries):
                scores = doc_emb @ q_emb[qi]
                order = np.argsort(-scores)
                qid = normalize_variant_string(q_texts[qi])
                # Keep only docs whose variant_id matches the normalized query ID.
                ranked = [doc_ids[i] for i in order
                          if qid is not None and doc_ids[i] == qid]
                ranked_per_query.append(ranked)
            for k in KS:
                agg = {"wrong": [], "recall": [], "mrr": [], "ndcg": []}
                for q, ranked in zip(queries, ranked_per_query):
                    m = retrieval_metrics(ranked, q["variant_id"],
                                          q["partner_ids"], k)
                    agg["wrong"].append(m["wrong"])
                    agg["recall"].append(m["recall"])
                    agg["mrr"].append(m["mrr"])
                    agg["ndcg"].append(m["ndcg"])
                rows.append(dict(
                    model=model, format=fmt, k=k,
                    wrong_variant_at_k=round(float(np.mean(agg["wrong"])), 4),
                    recall_at_k=round(float(np.mean(agg["recall"])), 4),
                    mrr=round(float(np.mean(agg["mrr"])), 4),
                    ndcg_at_k=round(float(np.mean(agg["ndcg"])), 4),
                ))

    write_csv(ROOT / "id-guardrail-metrics.csv",
              ["model", "format", "k", "wrong_variant_at_k", "recall_at_k",
               "mrr", "ndcg_at_k"], rows)

    for r in rows:
        if r["format"] == "medium" and r["k"] == 5:
            print(f"  {r['model']:>20} (medium, k=5): "
                  f"wrong_variant@5={r['wrong_variant_at_k']}  "
                  f"recall@5={r['recall_at_k']}")


if __name__ == "__main__":
    main()
```
