## Supplementary code and data: benchmark, experiment scripts, and result files for "Cosine Similarity Conflates Clinically Distinct Cancer Variants: A Case for Typed-Graph Retrieval in Precision Oncology Decision Support": experiment_loo_sensitivity.py

```
#!/usr/bin/env python3
"""
experiment_loo_sensitivity.py — Experiment 2.8: leave-one-pair-out sensitivity.

Holds out each of the nine clinically distinct pairs in turn and recomputes
the conflation rate at tau = 0.95 (medium format) on the remaining eight,
for the two biomedically pre-trained encoders. Demonstrates that the headline
conflation rate is not an artifact of any single pair.

Output:
  loo-sensitivity.csv   held_out_pair, model, conflation_rate_remaining_pairs

No embedding models are loaded; reads results.json. Runtime: < 2 seconds.
"""
from __future__ import annotations

from experiment_common import ROOT, BIOMEDICAL_MODELS, load_results, write_csv

TAU = 0.95
FORMAT = "medium"


def main() -> None:
    res = load_results()
    pairs = [p["pair_id"] for p in res["distinct_pairs"]]
    sims = res["similarities_distinct"]

    rows = []
    for held_out in pairs:
        remaining = [pid for pid in pairs if pid != held_out]
        for model in BIOMEDICAL_MODELS:
            k = sum(1 for pid in remaining if sims[model][FORMAT][pid] >= TAU)
            rows.append(dict(held_out_pair=held_out, model=model,
                             conflation_rate_remaining_pairs=round(k / len(remaining), 4)))

    write_csv(ROOT / "loo-sensitivity.csv",
              ["held_out_pair", "model", "conflation_rate_remaining_pairs"], rows)

    for model in BIOMEDICAL_MODELS:
        vals = [r["conflation_rate_remaining_pairs"] for r in rows if r["model"] == model]
        print(f"  {model} (medium, tau={TAU}): leave-one-out rates "
              f"min={min(vals)} max={max(vals)} over 9 held-out folds")


if __name__ == "__main__":
    main()
```
