## Supplementary code and data: benchmark, experiment scripts, and result files for "Cosine Similarity Conflates Clinically Distinct Cancer Variants: A Case for Typed-Graph Retrieval in Precision Oncology Decision Support": experiment_negative_controls.py

```
#!/usr/bin/env python3
"""
experiment_negative_controls.py — Experiment 2.2: template-matched negatives.

Without unrelated, template-matched controls a high cosine similarity could
be a template artifact ("<gene> <variant> in <tumor>" strings are all
similar) rather than true variant conflation. This script builds two
negative-control sets and scores them exactly as the main benchmark:

  Set A (cross-gene unrelated): each main-set variant_a is paired with a
    canonical driver variant from a different gene and a different tumor type
    (nine pairs). Template-matched, biologically unrelated.

  Set B (same-gene, non-canonical tumor): each distinct main-set variant_a is
    paired with itself in a non-canonical tumor type (eight pairs; KRAS G12C
    is variant_a in two main pairs, so eight distinct queries). Probes
    tumor-type sensitivity.

Outputs:
  negative-controls-cross-gene.csv             pair_id, model, format, cosine_similarity
  negative-controls-same-gene-other-tumor.csv  pair_id, model, format, cosine_similarity
  negative-controls-summary.csv                set, model, format, tau_* conflation rates

Runtime: a few minutes (loads up to three embedding models on first run).
"""
from __future__ import annotations

from experiment_common import (ROOT, MODELS, THRESHOLDS, DECOYS_SET_A,
                                SAME_GENE_OTHER_TUMOR, VARIANTS, embed, cosine,
                                load_pairs, variant_key, write_csv)

FORMATS = ("short", "medium", "long")


def render_main(v: dict, fmt: str) -> str:
    if fmt == "short":
        return v["short"]
    if fmt == "medium":
        return f"{v['short']} in {v['tumor']}"
    return v["asym"]


def render_decoy(d: dict, fmt: str) -> str:
    short = f"{d['gene']} {d['variant']}"
    if fmt == "short":
        return short
    if fmt == "medium":
        return f"{short} in {d['tumor']}"
    return d["context"]


def render_other_tumor(v: dict, tumor: str, fmt: str) -> str:
    if fmt == "short":
        return v["short"]
    if fmt == "medium":
        return f"{v['short']} in {tumor}"
    clause = v["asym"].split("; ", 1)[1]
    return f"{v['short']} in {tumor}; {clause}"


def conflation_summary(set_name: str, sims: dict[str, dict[str, list[float]]]) -> list[dict]:
    rows = []
    for model in MODELS:
        for fmt in FORMATS:
            vals = sims[model][fmt]
            row = dict(set=set_name, model=model, format=fmt)
            for tau in THRESHOLDS:
                row[f"tau_{tau}"] = round(sum(1 for s in vals if s >= tau) / len(vals), 4)
            rows.append(row)
    return rows


def main() -> None:
    distinct, _ = load_pairs()
    tau_cols = [f"tau_{t}" for t in THRESHOLDS]

    # ---- Set A: cross-gene unrelated -------------------------------------
    main_keys = [variant_key(p.gene, p.variant_a) for p in distinct]
    set_a = list(zip(main_keys, DECOYS_SET_A))  # 9 pairs

    a_rows = []
    a_sims: dict[str, dict[str, list[float]]] = {m: {f: [] for f in FORMATS} for m in MODELS}
    for model in MODELS:
        for fmt in FORMATS:
            texts_a = [render_main(VARIANTS[k], fmt) for k, _ in set_a]
            texts_b = [render_decoy(d, fmt) for _, d in set_a]
            ea, eb = embed(model, texts_a), embed(model, texts_b)
            for i, (va, vb) in enumerate(zip(ea, eb), start=1):
                s = cosine(va, vb)
                a_sims[model][fmt].append(s)
                a_rows.append(dict(pair_id=f"A{i}", model=model, format=fmt,
                                   cosine_similarity=round(s, 4)))
    write_csv(ROOT / "negative-controls-cross-gene.csv",
              ["pair_id", "model", "format", "cosine_similarity"], a_rows)

    # ---- Set B: same gene, non-canonical tumor ---------------------------
    set_b = list(SAME_GENE_OTHER_TUMOR.items())  # 8 (key, other_tumor)
    b_rows = []
    b_sims: dict[str, dict[str, list[float]]] = {m: {f: [] for f in FORMATS} for m in MODELS}
    for model in MODELS:
        for fmt in FORMATS:
            texts_1, texts_2 = [], []
            for key, other in set_b:
                v = VARIANTS[key]
                texts_1.append(render_main(v, fmt))
                texts_2.append(render_other_tumor(v, other, fmt))
            e1, e2 = embed(model, texts_1), embed(model, texts_2)
            for i, (v1, v2) in enumerate(zip(e1, e2), start=1):
                s = cosine(v1, v2)
                b_sims[model][fmt].append(s)
                b_rows.append(dict(pair_id=f"B{i}", model=model, format=fmt,
                                   cosine_similarity=round(s, 4)))
    write_csv(ROOT / "negative-controls-same-gene-other-tumor.csv",
              ["pair_id", "model", "format", "cosine_similarity"], b_rows)

    # ---- Summary ---------------------------------------------------------
    summary = (conflation_summary("cross-gene", a_sims)
               + conflation_summary("same-gene-other-tumor", b_sims))
    write_csv(ROOT / "negative-controls-summary.csv",
              ["set", "model", "format"] + tau_cols, summary)

    for r in summary:
        if r["format"] == "medium":
            print(f"  {r['set']:>22} {r['model']:>20} (medium): "
                  + "  ".join(f"{c}={r[c]}" for c in tau_cols))


if __name__ == "__main__":
    main()
```
