## Supplementary code and data: benchmark, experiment scripts, and result files for "Cosine Similarity Conflates Clinically Distinct Cancer Variants: A Case for Typed-Graph Retrieval in Precision Oncology Decision Support": experiment_separation.py

```
#!/usr/bin/env python3
"""
experiment_separation.py — Experiment 2.3: separation metrics.

Treats the equivalent-notation pairs as positives (label 1, "these strings
genuinely denote the same variant") and the clinically distinct pairs as
negatives (label 0, "these strings denote different variants"). For each
encoder and text format it asks whether a cosine threshold can separate the
two classes, with:

  * ROC-AUC (+ bootstrap 95% CI, 10,000 resamples)
  * PR-AUC (average precision)
  * Kolmogorov-Smirnov statistic and p-value
  * Cliff's delta (positives vs negatives)
  * Youden's J optimal cosine threshold

An AUC near 0.5 means equivalent-notation and clinically distinct pairs are
not separable by cosine; an AUC below 0.5 means clinically distinct pairs
score even higher than known-equivalent notations.

Output:
  separation-metrics.csv

No embedding models are loaded; reads results.json. Runtime: ~10 seconds.
"""
from __future__ import annotations

import numpy as np
from scipy.stats import ks_2samp
from sklearn.metrics import average_precision_score, roc_auc_score, roc_curve

from experiment_common import ROOT, MODELS, load_results, write_csv

RNG = np.random.default_rng(20260505)
N_BOOT = 10_000


def cliffs_delta(pos: np.ndarray, neg: np.ndarray) -> float:
    gt = sum((p > neg).sum() for p in pos)
    lt = sum((p < neg).sum() for p in pos)
    return (gt - lt) / (len(pos) * len(neg))


def auc_with_label(labels: np.ndarray, scores: np.ndarray) -> float:
    """ROC-AUC oriented so that label 1 = positive (equivalent-notation)."""
    return roc_auc_score(labels, scores)


def main() -> None:
    res = load_results()
    formats = res["formats"]
    pairs = [p["pair_id"] for p in res["distinct_pairs"]]
    eq_ids = [p["pair_id"] for p in res["equivalent_pairs"]]
    sims_d = res["similarities_distinct"]
    sims_e = res["similarities_equivalent"]

    rows = []
    for model in MODELS:
        pos = np.array([sims_e[model][e] for e in eq_ids], dtype=float)
        for fmt in list(formats) + ["averaged"]:
            if fmt == "averaged":
                neg = np.array([np.mean([sims_d[model][f][pid] for f in formats])
                                for pid in pairs], dtype=float)
            else:
                neg = np.array([sims_d[model][fmt][pid] for pid in pairs], dtype=float)

            labels = np.array([1] * len(pos) + [0] * len(neg))
            scores = np.concatenate([pos, neg])

            roc_auc = auc_with_label(labels, scores)
            pr_auc = average_precision_score(labels, scores)
            ks = ks_2samp(pos, neg)
            delta = cliffs_delta(pos, neg)

            fpr, tpr, thr = roc_curve(labels, scores)
            j = tpr - fpr
            youden_thr = float(thr[int(np.argmax(j))])

            # Stratified bootstrap CI for ROC-AUC.
            n_pos, n_neg = len(pos), len(neg)
            boot = np.empty(N_BOOT)
            for i in range(N_BOOT):
                pi = RNG.integers(0, n_pos, n_pos)
                ni = RNG.integers(0, n_neg, n_neg)
                bl = np.array([1] * n_pos + [0] * n_neg)
                bs = np.concatenate([pos[pi], neg[ni]])
                boot[i] = roc_auc_score(bl, bs)
            ci_lo, ci_hi = np.percentile(boot, [2.5, 97.5])

            rows.append(dict(
                model=model, format=fmt,
                roc_auc=round(float(roc_auc), 4),
                roc_auc_ci_lower=round(float(ci_lo), 4),
                roc_auc_ci_upper=round(float(ci_hi), 4),
                pr_auc=round(float(pr_auc), 4),
                ks_stat=round(float(ks.statistic), 4),
                ks_p=round(float(ks.pvalue), 4),
                cliffs_delta=round(float(delta), 4),
                youden_j_threshold=round(youden_thr, 4),
            ))

    write_csv(ROOT / "separation-metrics.csv",
              ["model", "format", "roc_auc", "roc_auc_ci_lower", "roc_auc_ci_upper",
               "pr_auc", "ks_stat", "ks_p", "cliffs_delta", "youden_j_threshold"], rows)

    for r in rows:
        if r["format"] == "medium":
            print(f"  {r['model']:>20} (medium): ROC-AUC={r['roc_auc']} "
                  f"[{r['roc_auc_ci_lower']}, {r['roc_auc_ci_upper']}]  "
                  f"Cliff's delta={r['cliffs_delta']}")


if __name__ == "__main__":
    main()
```
