## Supplementary code and data: benchmark, experiment scripts, and result files for "Cosine Similarity Conflates Clinically Distinct Cancer Variants: A Case for Typed-Graph Retrieval in Precision Oncology Decision Support": experiment_typed_graph.py

```
#!/usr/bin/env python3
"""
experiment_typed_graph.py — Experiment 2.7: end-to-end typed-graph pipeline.

Builds a minimal in-memory typed property graph (networkx, so no Neo4j
dependency) with Gene, Variant, Drug, and EvidenceStatement nodes and
INDICATES_RESPONSE_TO / CONTRAINDICATES / DESCRIBED_BY edges, populated from
the sixteen distinct variants of the nine-pair benchmark.

It then runs the full ingestion pipeline on every benchmark variant string in
all three text formats:
  ingest -> normalize_variant_string() -> exact graph lookup -> drug retrieval

and measures normalization accuracy, drug-retrieval accuracy, and the
wrong-variant retrieval rate. Normalization uses general regex rules (shared
in experiment_common.py); they are not hand-tuned to the benchmark, so
non-point-mutation classes are characterized honestly.

Outputs:
  typed-graph-results.csv   variant, format, normalized_correctly,
                            drug_set_retrieved, wrong_variant_retrieved
  typed-graph-summary.csv   aggregate accuracy metrics, per format and overall

No embedding models are loaded. Runtime: < 2 seconds.
"""
from __future__ import annotations

import networkx as nx

from experiment_common import ROOT, VARIANTS, normalize_variant_string, write_csv

FORMATS = ("short", "medium", "long")


def build_graph() -> tuple[nx.DiGraph, dict[tuple[str, str], str]]:
    """Return the typed graph and a {(gene, hgvs_protein): variant_node_id} index."""
    g = nx.DiGraph()
    index: dict[tuple[str, str], str] = {}
    for key, v in VARIANTS.items():
        gene_id = v["gene"]
        var_id = f"{v['gene']}:{v['hgvs_protein']}"
        ev_id = f"EV:{var_id}"
        g.add_node(gene_id, ntype="Gene", symbol=v["gene"])
        g.add_node(var_id, ntype="Variant", gene=v["gene"],
                   hgvs_protein=v["hgvs_protein"], tier=v["tier"], label=key)
        g.add_node(ev_id, ntype="EvidenceStatement", text=v["asym"],
                   amp_asco_cap_tier=v["tier"])
        g.add_edge(var_id, gene_id, etype="OF_GENE")
        g.add_edge(var_id, ev_id, etype="DESCRIBED_BY")
        for drug in v["drugs"]:
            g.add_node(drug, ntype="Drug", name=drug)
            g.add_edge(var_id, drug, etype="INDICATES_RESPONSE_TO",
                       evidence_level=v["tier"], source="benchmark")
        index[(v["gene"], v["hgvs_protein"])] = var_id
    return g, index


def drugs_of(g: nx.DiGraph, var_id: str) -> list[str]:
    return sorted(t for _, t, d in g.out_edges(var_id, data=True)
                  if d.get("etype") == "INDICATES_RESPONSE_TO")


def render(v: dict, fmt: str) -> str:
    if fmt == "short":
        return v["short"]
    if fmt == "medium":
        return f"{v['short']} in {v['tumor']}"
    return v["asym"]  # long: full variant-specific clinical sentence


def main() -> None:
    g, index = build_graph()
    n_var = sum(1 for _, d in g.nodes(data=True) if d["ntype"] == "Variant")
    n_drug = sum(1 for _, d in g.nodes(data=True) if d["ntype"] == "Drug")
    print(f"Typed graph: {n_var} Variant nodes, {n_drug} Drug nodes, "
          f"{g.number_of_edges()} typed edges.")

    rows = []
    for key, v in VARIANTS.items():
        intended_id = index[(v["gene"], v["hgvs_protein"])]
        intended_drugs = drugs_of(g, intended_id)
        for fmt in FORMATS:
            text = render(v, fmt)
            norm = normalize_variant_string(text)

            normalized_correctly = norm == (v["gene"], v["hgvs_protein"])
            retrieved_id = index.get(norm) if norm else None
            wrong_variant = retrieved_id is not None and retrieved_id != intended_id

            if retrieved_id is None:
                drug_set = "UNRESOLVED"
            else:
                drugs = drugs_of(g, retrieved_id)
                drug_set = "; ".join(drugs) if drugs else "none (Tier III)"

            rows.append(dict(
                variant=key, format=fmt,
                normalized_correctly=normalized_correctly,
                drug_set_retrieved=drug_set,
                wrong_variant_retrieved=wrong_variant,
                _correct_drugs=drug_set == ("; ".join(intended_drugs)
                                            if intended_drugs else "none (Tier III)"),
            ))

    write_csv(ROOT / "typed-graph-results.csv",
              ["variant", "format", "normalized_correctly", "drug_set_retrieved",
               "wrong_variant_retrieved"],
              [{k: r[k] for k in ("variant", "format", "normalized_correctly",
                                  "drug_set_retrieved", "wrong_variant_retrieved")}
               for r in rows])

    # Aggregate summary, per format and overall.
    summary = []
    for fmt in list(FORMATS) + ["all"]:
        sub = [r for r in rows if fmt == "all" or r["format"] == fmt]
        n = len(sub)
        n_norm = sum(r["normalized_correctly"] for r in sub)
        n_drug_ok = sum(r["_correct_drugs"] for r in sub if r["normalized_correctly"])
        n_wrong = sum(r["wrong_variant_retrieved"] for r in sub)
        summary.append(dict(
            format=fmt, n=n,
            normalization_accuracy=round(n_norm / n, 4),
            drug_retrieval_accuracy=round(n_drug_ok / n_norm, 4) if n_norm else 0.0,
            wrong_variant_retrieval_rate=round(n_wrong / n, 4),
        ))
    write_csv(ROOT / "typed-graph-summary.csv",
              ["format", "n", "normalization_accuracy", "drug_retrieval_accuracy",
               "wrong_variant_retrieval_rate"], summary)

    overall = summary[-1]
    print(f"  normalization accuracy   : {overall['normalization_accuracy']}")
    print(f"  drug-retrieval accuracy  : {overall['drug_retrieval_accuracy']} "
          f"(of correctly normalized)")
    print(f"  wrong-variant retrieval  : {overall['wrong_variant_retrieval_rate']}")
    failures = [r["variant"] for r in rows if not r["normalized_correctly"]]
    if failures:
        print(f"  normalization failures   : {sorted(set(failures))}")


if __name__ == "__main__":
    main()
```
