## Supplementary code and data: benchmark, experiment scripts, and result files for "Cosine Similarity Conflates Clinically Distinct Cancer Variants: A Case for Typed-Graph Retrieval in Precision Oncology Decision Support": README.md

# Variant-Pair Conflation Benchmark — Reproducibility Guide (v2)

This directory holds the manuscript *"Cosine Similarity Conflates Clinically
Distinct Cancer Variants: A Case for Typed-Graph Retrieval in Precision
Oncology Decision Support"* (Khan, U., UNMIRI LLC) and the runnable
experiments that produce its results.

- **v1 preprint:** bioRxiv, doi:10.64898/2026.05.05.723102.
- **v2 (this revision):** adds eight experiments in response to an external
  methodological critique, and is prepared for journal submission (target:
  *JAMIA*; alternative: *NEJM AI*).

Everything runs on a single laptop, CPU-only, with **no external API
dependency**. The only network access is the one-time download of three
open-source embedding models from the Hugging Face Hub.

## Files in this directory

### Manuscript
| File | What it is |
|---|---|
| `manuscript.md` | The paper (source). |
| `manuscript.docx` | Word render. Regenerated from the markdown; do not hand-edit. |
| `figure-1.png` | Figure 1, the per-pair similarity heatmap. |
| `v2-revision-summary.md` | Summary of what changed in the v2 revision. |

### Benchmark data
| File | What it is |
|---|---|
| `variant-pairs.csv` | The 9 clinically distinct variant pairs (curated from CIViC). |
| `equivalent-pairs.csv` | 6 equivalent-notation pairs (positive control). |

### Code
| File | What it is |
|---|---|
| `experiment.py` | v1 benchmark: pairwise cosine similarity, three encoders, three formats. |
| `experiment_common.py` | Shared helpers for the v2 experiments: embedding cache, the canonical 16-variant table, negative-control and decoy sets, the variant-string normalizer, and the retrieval corpus. |
| `experiment_ci.py` | **2.1** Exact binomial 95% CIs + bootstrap CIs. |
| `experiment_negative_controls.py` | **2.2** Template-matched negative controls (cross-gene; same-gene other-tumor). |
| `experiment_separation.py` | **2.3** ROC-AUC / PR-AUC / KS / Cliff's delta separation metrics. |
| `experiment_asymmetric_long_format.py` | **2.4** Variant-specific (asymmetric) long format + Jaccard overlap. |
| `experiment_corpus_retrieval.py` | **2.5** Corpus-level ranked retrieval (52-doc corpus). |
| `experiment_id_guardrail.py` | **2.6** Cosine + exact variant-ID guardrail baseline. |
| `experiment_typed_graph.py` | **2.7** End-to-end typed-graph pipeline (networkx). |
| `experiment_loo_sensitivity.py` | **2.8** Leave-one-pair-out sensitivity. |
| `requirements.txt` | Python dependencies. |

## Step 1 — Set up

```bash
cd docs/papers/graphrag-vs-vector-rag-2026
python3.12 -m venv .venv
source .venv/bin/activate
pip install -r requirements.txt
```

First run downloads three embedding models (~3 GB total: PubMedBERT ~440 MB,
MedCPT ~440 MB, BGE-large-en-v1.5 ~1.3 GB), cached afterward under
`~/.cache/huggingface`.

## Step 2 — Run the experiments

```bash
python3 experiment.py                          # v1 benchmark
python3 experiment_ci.py                        # 2.1
python3 experiment_negative_controls.py         # 2.2
python3 experiment_separation.py                # 2.3
python3 experiment_asymmetric_long_format.py    # 2.4
python3 experiment_corpus_retrieval.py          # 2.5
python3 experiment_id_guardrail.py              # 2.6  (run after 2.5)
python3 experiment_typed_graph.py               # 2.7
python3 experiment_loo_sensitivity.py           # 2.8
```

Each script is self-contained and writes its own output files; the scripts
may be run in any order, except that `experiment_ci.py`,
`experiment_separation.py`, and `experiment_loo_sensitivity.py` read
`results.json` (so run `experiment.py` first), and `experiment_id_guardrail.py`
reuses the cached corpus embeddings from `experiment_corpus_retrieval.py`.

**Runtime.** With the models already downloaded, the full v1 + v2 suite
re-runs in **under 30 minutes** on a laptop CPU (most of that is model
loading; the embedding computations themselves are small). Embeddings are
cached on disk in `.emb-cache/` so the v2 scripts share computation and a
re-run is near-instant. Delete `.emb-cache/` to force a clean recomputation.
`experiment_ci.py`, `experiment_separation.py`, `experiment_typed_graph.py`,
and `experiment_loo_sensitivity.py` load no embedding model at all.

## Output files and schemas

### v1 outputs
| File | Schema / contents |
|---|---|
| `results.json` | Full per-pair, per-model, per-format similarity scores. |
| `conflation-rates.csv` | `model, tau_0.85, tau_0.9, tau_0.95, tau_0.99` (Table 1). |
| `format-rates.csv` | `format, PubMedBERT, MedCPT, BGE-large-en-v1.5` at τ=0.95 (Table 2). |
| `positive-control.json` | Per-encoder mean/sd/min/max for equivalent-form pairs. |
| `figure-1.png` | Per-pair medium-format heatmap (Figure 1). |

### v2 outputs
| File | Schema / contents |
|---|---|
| `conflation-rates-with-ci.csv` | `model, threshold, format, n, k, proportion, ci_lower, ci_upper` — exact binomial 95% CIs. |
| `positive-control-with-ci.json` | Per-encoder `n, mean, sd, ci_lower, ci_upper, n_bootstrap` (10,000 bootstrap resamples). |
| `negative-controls-cross-gene.csv` | `pair_id, model, format, cosine_similarity` — Set A (cross-gene unrelated). |
| `negative-controls-same-gene-other-tumor.csv` | `pair_id, model, format, cosine_similarity` — Set B. |
| `negative-controls-summary.csv` | `set, model, format, tau_0.85, tau_0.9, tau_0.95, tau_0.99`. |
| `separation-metrics.csv` | `model, format, roc_auc, roc_auc_ci_lower, roc_auc_ci_upper, pr_auc, ks_stat, ks_p, cliffs_delta, youden_j_threshold`. |
| `asymmetric-long-format-rates.csv` | `format, PubMedBERT, MedCPT, BGE-large-en-v1.5` — conflation rate per threshold (same column schema as `format-rates.csv`). |
| `asymmetric-long-per-pair.csv` | `pair_id, model, cosine_similarity`. |
| `asymmetric-long-jaccard.csv` | `pair_id, gene, jaccard, shared_tokens, union_tokens`. |
| `corpus-retrieval-metrics.csv` | `model, format, k, wrong_variant_at_k, recall_at_k, mrr, ndcg_at_k`. |
| `id-guardrail-metrics.csv` | Same schema as `corpus-retrieval-metrics.csv`. |
| `typed-graph-results.csv` | `variant, format, normalized_correctly, drug_set_retrieved, wrong_variant_retrieved`. |
| `typed-graph-summary.csv` | `format, n, normalization_accuracy, drug_retrieval_accuracy, wrong_variant_retrieval_rate`. |
| `loo-sensitivity.csv` | `held_out_pair, model, conflation_rate_remaining_pairs`. |

All v1 output formats are unchanged in v2; v2 adds new files rather than
restructuring existing ones.

## Dependencies

`requirements.txt` pins the v1 stack (`sentence-transformers`,
`transformers`, `torch`, `numpy`, `matplotlib`) and adds three packages for
the v2 experiments: **`scipy`** (exact binomial CIs, Kolmogorov-Smirnov
test), **`scikit-learn`** (ROC-AUC, PR-AUC), and **`networkx`** (the
in-memory typed property graph in Experiment 2.7). No other dependency is
introduced; the embedding cache and the typed graph are both kept in-process
so the suite needs no database and no network API.

## Reproducibility notes

- All randomized procedures (bootstrap CIs) use a fixed seed (`20260505`), so
  reported intervals are reproducible to the digit.
- Embeddings are deterministic given (model, text); `.emb-cache/` only
  accelerates re-runs and never changes a result.
- `experiment.py` re-run reproduces `results.json`, `conflation-rates.csv`,
  and `format-rates.csv` exactly.

## Citation

> Khan, U. *Cosine Similarity Conflates Clinically Distinct Cancer Variants:
> A Case for Typed-Graph Retrieval in Precision Oncology Decision Support.*
> v2, 2026. bioRxiv doi:10.64898/2026.05.05.723102. Benchmark and code:
> Zenodo doi:10.5281/zenodo.20042352; github.com/unmirihealth/unmiri-ngs-fhir-schema.

Benchmark and code are licensed CC BY 4.0 (data) / Apache 2.0 (schema) to
enable replication and extension.
